# Distinct neural representational geometries of numerosity in early visual and association regions of dorsal and ventral streams

**DOI:** 10.1101/2023.12.18.571155

**Authors:** Alireza Karami, Elisa Castaldi, Evelyn Eger, Manuela Piazza

**Author notes:** **Corresponding authors email address:** Alireza Karami, Manuela Piazza.

## Abstract

Visual numerosity, a feature of visual sets traditionally associated with the parietal cortex, has recently been suggested to be encoded in a broader range of cortical regions, including early visual and associative cortices of both streams. However, a clear picture of how numerosity information is represented alongside other non-numeric features across the cortex is still lacking. To address this, we conducted a whole-brain 3T fMRI study with thirty-one adult subjects (eighteen females) engaged in a numerosity estimation task involving visual sets varying in number, item size, total item area, total field area, and density, thus allowing us to perform a tight control of the stimulus properties. Using model-based semipartial correlation representational similarity analyses, we found that numerosity is represented independently of other visual properties already in early visual areas and amplified in both retinotopic and non-retinotopic associative areas along both the dorsal and ventral streams. Dimensionality reduction analyses of BOLD activity patterns revealed clearly distinct neural representational geometries of numerosity across brain regions: a one-dimensional representation of numerical rank in early visual and in associative retinotopic regions of the ventral stream, and a curved structure encoding rank-order together with distance to end-point in associative regions of both the dorsal and ventral stream. These findings indicate that numerosity is represented over and above other visual features broadly in the brain but with substantially different neural coding schemes across regions, and are discussed in light of two non necessarily exclusive interpretations.

## Introduction

The ability to estimate the number of objects (numerosity) in the environment is ontogenetically precocious and phylogenetically ancient. In animals it is of high adaptive value (Nieder, 2020). In humans it was suggested to also play a role in scaffolding the acquisitions of symbolic numeracy (Piazza, 2010; Halberda et al., 2008; Decarli et al., 2023), representing a potential target for math education and remediation. Single cell recording studies indicate that in both in the animal (e.g., Nieder & Miller, 2003; Wagener et al., 2018; Kobylkov et al., 2022) and the human brain (Kutter et al., 2018) there are numerosity-tuned neurons, whose Gaussian tuning functions reflect scalar variability and underlie behavior in numerosity comparison and matching tasks, similarly adhering to Weber’s law (Gallistel & Gelman, 1992; Merten & Nieder, 2009; Ditz & Nieder, 2016; Piantadosi & Cantlon, 2017). This approximate and compressed code can also be inferred from the population-level responses to numerosity as measured by fMRI using multivariate pattern analyses, adaptation, or population receptive field mapping approaches (e.g., Piazza et al., 2004; Eger et al., 2009; Harvey et al., 2013). However, because numerosity is necessarily coupled with other visual characteristics of the sets (e.g., more items tend to occupy a larger area, or to be more densely spaced), establishing the degree to which the observed neural response to numerosity is distinct from the response to other visual attributes is not trivial. Castaldi et al. (2019) measured the human fMRI BOLD response to numerosity and approached this issue combining multivariate representational similarity analyses with multiple regression, estimating the brain activity evoked by numerosity once taking into account the effects of other relevant non-numerical variables at the same time (Castaldi et al., 2019). This study, solely focussing on the dorsal stream, demonstrated that numerosity is represented over and above other visual features across all retinotopic regions along the dorsal stream, and, especially when task relevant, amplified in parietal areas. In terms of localization, these results seemed in agreement with single cell recordings in macaques (Nieder & Miller, 2003; Nieder & Miller, 2004) and fMRI studies in humans (e.g., Piazza et al., 2004; Cantlon et al., 2006; Castelli et al., 2006; Ansari & Dhital, 2006) that pointed to the parietal cortex as the key brain region for numerosity processing (see for a review: Eger, 2016; Faye et al., 2019). They were also consistent with the neuropsychological literature that associates deficits in numerosity processing to parietal cortex damage (Warrington & James, 1967; Lemer et al., 2003). Partially biased by these initial observations, several later key fMRI studies on numerosity restricted the brain data acquisition to a limited volume centered on parietal cortex (e.g. Eger et al., 2009; Castaldi et al., 2019) or focussed the analyses on parietal cortex using an ROI approach (e.g., Bulthè et al., 2015; Castaldi et al., 2020). While a few studies looking at the whole brain did sometimes report numerosity-related response outside parietal cortex, both in the frontal and in the occipito-temporal cortex (e.g., Bulthè et al., 2014; Harvey & Dumoulin, 2017), they mostly tested small numerosities, which some suggest to be elaborated by a dedicated mechanism, referred to as “subitizing”, potentially different from the one at play with large numerosities (Revkin et al., 2008; Kutter et al., 2023). One notable exception is the recent work of Cai et al. (2021), who employed a population receptive field mapping (pRF) method (Harvey & Dumoulin, 2017) and found that the numerotopic maps that encode small numerosities in association regions in the dorsal and ventral stream also encode large numerosities. Two important limitations from this study, however, call for further confirmations of this stand-alone report:

1. In the study non-numerical features were neither controlled for nor their impact on brain response analyzed: in all trials total surface area was held constant across numerosities, thus as a consequence both dot size and density were 100% correlated with number. This is particularly problematic as the authors themselves had previously shown that numerosity and object size are represented in overlapping topographic maps (Harvey et al., 2015).
2. They presented stimuli in strictly ordered sequences (increasing or decreasing numerosity). This approach, while ideal for pRF modeling, likely creates expectations (e.g. Puri et al., 2009; Esterman & Yantis, 2009; for a review see: Summerfield & De Lange, 2014) and attentional biases (e.g. Jehee et al., 2011; Ester et al., 2016; Lage-Castellanos et al., 2022), the effect of which cannot be readily disentangled from the effect of numerosity itself. Thus, the question of whether large numerosity is encoded over and above the other visual properties solely along the dorsal stream, or whether it is also represented in other associative regions of the human brain still remains open.

Another set of studies that found numerosity information outside the parietal cortex are the ones mainly performed by Fornciai and collaborators (Fornaciai et al., 2017; Fornaciai & Park, 2018) who used EEG and fMRI and found pure numerosity information over midline occipital electrodes very early in the time course, suggesting that it is initially extracted in early visual regions (V1 / V2). Early visual regions involvement in encoding numerosity was also previously reported by Lasne et al. (2019), who scanned subjects with fMRI a reduced brain volume including early visual and parietal sites and showed that numerosity could be successfully decoded not only in parietal but also in V1. Interestingly, despite the various differences in stimuli, tasks, and analytical approaches, all previous studies on numerosity that employed model-based methods—such as decoding, RSA, or PRF modeling (e.g., Eger et al., 2009; Harvey et al., 2013; Castaldi et al., 2019; Cai et al., 2021)—have assumed that numerosity is encoded in a unidimensional space (i.e., a line) of well-ordered magnitude, while to the best of our knowledge no one has tried to approach the data using model-free analytical approaches, which could be of great value in revealing coding schemes that were not previously predicted (e.g., Luyckx et al., 2019; Nelli et al., 2023). This model-free approach could help in resolving the question of whether the geometry of the neural representation of numerosity varies in different brain regions, and, if so, how.

In order to probe these questions, in the current study we recorded the BOLD signal from the whole brain of subjects looking at sets of different number of dots, and analyzed the data using both a model-based representational similarity analysis in pre-defined ROIs and across the whole brain and model-free dimensionality reduction technique to better characterize the neural representational geometries of numerosity across regions.

## Materials and Methods

### Participants

Thirty-seven healthy adults (twenty-two females; mean age 21.9 years) with normal or corrected vision participated in the study. The ethics committee of the University of Trento (Italy) approved the study, and all participants gave written informed consent and were reimbursed for their time. Given their excessive head motion (translation, in one of the directions x,y or z, greater than 3mm, or rotation, around one of the axes, greater than 2 degrees) or poor behavioral performance (accuracy < 65%), data from six participants (four for poor behavioral performance and two for excessive head motion) were excluded from the final analysis. This led to the final sample of thirty-one subjects (eighteen females; mean age 21.9 years).

### Experimental Design and Statistical Analyses

#### Stimuli and procedure

Participants were familiarized with the task by practicing 20 trials outside of the MRI before the experiment. During fMRI scanning, arrays of black dots on a mid-gray background were centrally shown to participants. Dots orthogonally varied in number, average item area, and total field area (similar to Castaldi et al., 2019).

There were 32 conditions (resulting from crossing 4 numerosities, 4 average item areas, and 2 total field areas): six, ten, seventeen, or twenty-nine dots were presented with varying average item area (0.04, 0.07, 0.12, 0.2 visual square degree) that were made to fit within a small or large total field area (defined by a virtual circle of either about 9 or 13.5 visual degree diameter; Figure 1.A). Numbers and average item areas were chosen based on previous behavioral studies (Castaldi et al., 2018; Castaldi et al., 2019) to be equally discriminable. Total field areas were selected to have suitably sparse arrays of dots (1 dot/vd^2^) to be within the numerosity estimation and not the density estimation regime (Anobile et al., 2013).

**Figure 1.**
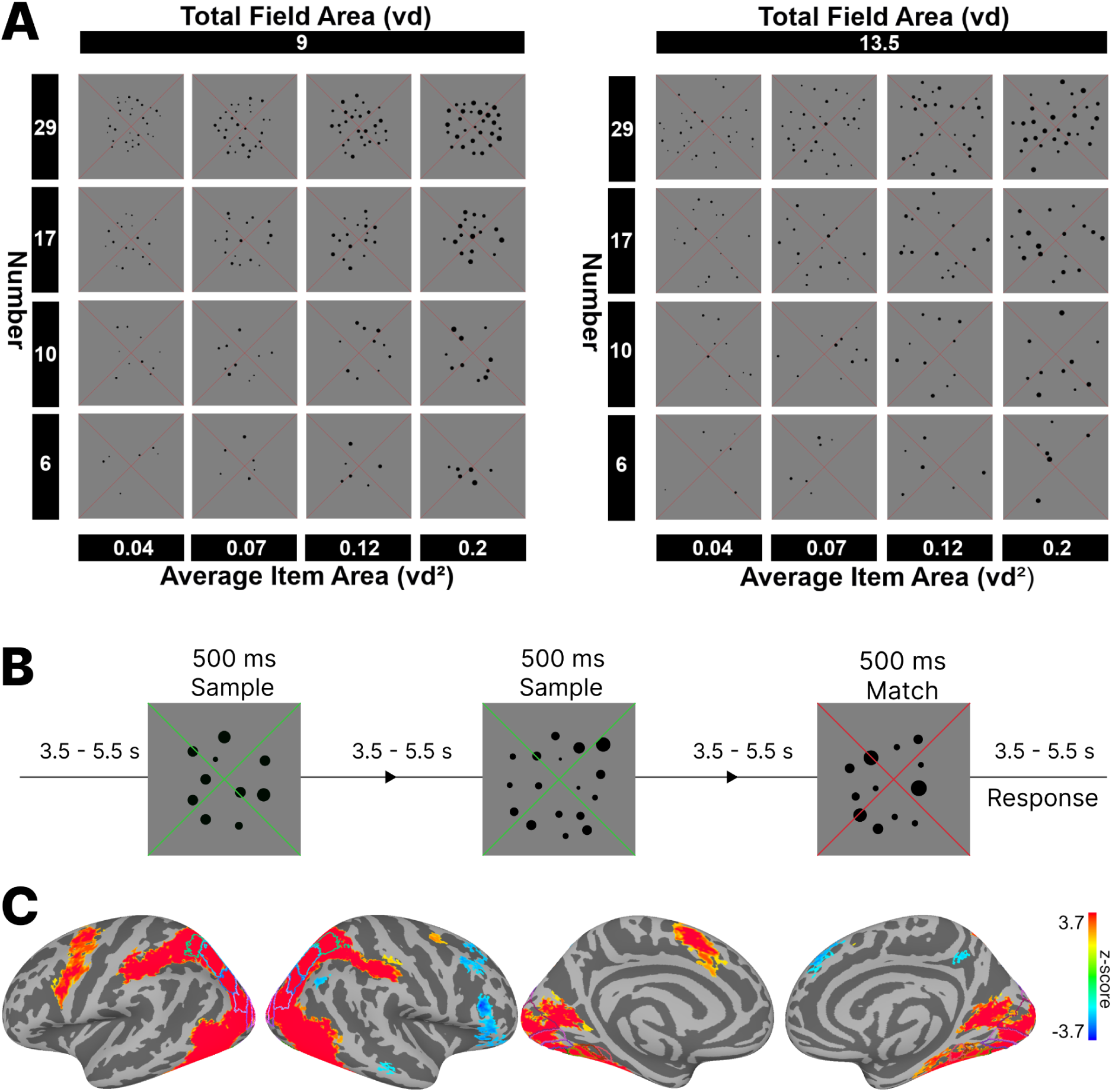
Stimulus set, design, and univariate effect of stimuli (A) Illustration of the stimulus conditions. Stimuli orthogonally varied in number (6,10,17,29), average item area (0.04, 0.07, 0.12 and 0.2 visual square degree), and total field area (9 and 13.5 visual degree). (B) Illustration of the temporal presentation of the stimuli during scanning. Participants were instructed to attend to the number of dots and keep the number in mind until the next set of dots was shown (after a variable time interval of 3.5–5.5 s). From time to time, the color of the fixation cross changed from red to green. When the color changed, subjects were required to compare the number of dots in the current set, (match stimulus), with the previous one (sample stimulus) by pressing a button. (C) Results obtained from the univariate surface-based group analysis (n = 31). The maps show the activation elicited for all sample stimuli contrasted against the implicit baseline. Activation maps are thresholded at p < 0.01, TFCE corrected, and displayed on Freesurfer’s fsaverage surface with colored outlines identifying ROIs along the early visual areas (V1, V2, V3), the dorsal (V3AB, V7, IPS12, IPS345), and the ventral stream (hV4, VO1, VO2, PHC1, PHC2) from a surface-based probabilistic atlas (Wang et al., 2015).

In each trial, a set of dots was presented for 500 ms over a wide thin red fixation cross, and participants were required to estimate their number and keep them in memory until, after a variable ISI of 3.5–5.5 s, the next set of dots appeared (Figure 1.B). When the color of the fixation cross changed from red to green, subjects were required to compare the number of dots in the current set (match stimulus) with the previous one and determine whether it was larger or smaller by pressing one of two buttons following the instructions. After a button press, the background cross turned red again, and a variable blank delay of 3.5–5.5 s preceded the following trial. Match stimuli were designed to be ∼2 JNDs larger or smaller in numerosity than the previous sample stimulus, based on an average numerosity Weber fraction estimated on an independent group of healthy adults (Castaldi et al., 2018), while the other dimensions (total field area and average item area) were the same. Match trials occurred ∼20% of the time.

The experiment consisted of six runs, with two blocks within each run. Each block consisted of thirty-six trials: four match trials and thirty-two sample trials, one for each condition (4 number × 4 average item area × 2 total field area). After the third run, in the middle of the experiment, participants’ hand-response correspondence was switched. There was a brief practice session at the beginning of the experiment and after changing the hand assignment. Each run lasted ∼7 minutes.

#### fMRI recordings and preprocessing

Functional images were acquired at the Center for Mind/Brain Sciences (CIMeC) with a SIEMENS MAGNETOM PRISMA 3T with gradient insert of 80mT/m max and using a SIEMENS Head/Neck 64-channel phased array coil. Visual stimuli were presented through a mirror system connected to a 42″ LCD monitor (MR-compatible, NordicNeuroLab) positioned at the back of the magnet bore. Functional images were acquired using echo-planar (EPI) T2*-weighted fat-saturation echo-planar image (EPI) volumes with 1.75 mm isotropic voxels using a multi-band sequence (Moeller et al., 2010) (https://www.cmrr.umn.edu/multiband/, multi-band [MB] = 3, GRAPPA acceleration with [IPAT] = 1, partial Fourier [PF] = 7/8, matrix = 120×120, repetition time [TR] = 2 s, echo time [TE] = 31.2 ms, echo spacing [ES] = 0.62 ms, flip angle [FA] = 60°, bandwidth [BW] = 2450 Hz/px, phase-encode direction Anterior >> Posterior). In total, 1206 volumes from the six experimental runs made up the functional acquisition. A whole-brain gradient echo B0 map, matched for spatial resolution, was acquired after the functional scans for fieldmap-based correction of susceptibility-induced geometric distortions. T1-weighted anatomical images were acquired at 1 mm isotropic resolution using an MPRAGE sequence (GRAPPA acceleration with [IPAT] = 2, matrix = 176×256, repetition time [TR] = 2530 s, echo time [TE] = 1.69 ms, time of inversion [TI] = 1100 ms, flip angle [FA] = 7°, bandwidth [BW] = 650 Hz/px). Padding and tape were used to reduce head movement. In their left and right hands, participants held two response buttons. Stimuli were presented using a custom-written Psychtoolbox 3 (Brainard, 1997) script running in MATLAB R2018 (The MathWorks, Inc., Natick, MA).

Functional images were preprocessed in MATLAB R2019 using the Statistical Parametric Mapping Software (SPM12, https://www.fil.ion.ucl.ac.uk/spm/software/spm12/). Preprocessing included the following steps: Slice-time correction of functional images to the middle slice, applying distortion correction to all functional images, realignment of each scan to the mean of each run, co-registration of the anatomical scan to the mean functional image, and segmentation of the anatomical image into native space tissue components.

The preprocessed EPI images (in subjects’ native space) were high-pass filtered at 128 s and pre-whitened by means of an autoregressive model AR(1). A general linear model (GLM) was used to estimate subject-specific beta weights. For each run, thirty-two regressors of interest were included for all sample stimuli (4 number × 4 average item area × 2 total field area). Regressors for match stimuli, left hand, and right hand were also included. Nuisance regressors were identified with The PhysIO toolbox (Kasper et al., 2017) using six motion parameters and CompCor with five components (Behzadi et al., 2007) and were included in the GLM.

The surface of each subject was generated using Freesurfer 6 (https://surfer.nmr.mgh.harvard.edu/) (Fischl, 2012). The surfaces were then converted to a SUMA standard mesh of 141,000 nodes per hemisphere (Saad et al., 2004) from each participant’s anatomical scan using algorithms implemented in the Surfing toolbox (https://surfing.sourceforge.net) (Oosterhof et al., 2011) to produce node-to-node anatomical correspondence across participants’ surfaces. For each subject, the parameter estimates (beta weights) for each of the 32 regressors of interest were converted into a t-statistic and projected on the SUMA standard mesh using AFNI’s 3dVol2Surf (https://afni.nimh.nih.gov/) (Cox, 1996) with the “average” mapping algorithm, which roughly represents the value at each vertex of the surface as the average value along a line connecting the smooth white matter and pial surfaces.

#### fMRI data analysis

First, in order to visualize the brain regions with activation during the stimuli, we smoothed with a Gaussian 4 mm FWHM filter using the SurfSmooth function with the HEAT_07 smoothing method the contrast map of sample stimulus against the implicit baseline (Chung et al., 2005). We then performed a surface-based random-effects group analysis using a one-sample t-test. The result was then corrected using threshold-free cluster enhancement (TFCE; Smith & Nichols, 2009) using Monte Carlo simulations with 10,000 permutations, as implemented in the CoSMoMVPA MATLAB toolbox (Oosterhof et al., 2016) and projected onto the fsaverage surface for visualization (thresholded at p < 0.01, one-tailed).

Second, in order to test if and in which brain regions the representations of numerical and the non-numerical features could be dissociated, we used Representational Similarity Analysis (RSA; Kriegeskorte et al., 2008; Kriegeskorte & Kievit, 2013) which allows to assess the effects of the experimental conditions on distributed activity patterns. We adopted two approaches: a Region Of Interest (ROI) and a whole brain searchlight approach (see below). In both cases we used the t-statistics from the first-level analysis to extract the neural representational dissimilarity matrix (RDM) by computing the correlation distance (Pearson correlation) between activation patterns for each pair of conditions. Each pattern underwent voxel-wise scaling by demeaning the data across conditions before computing the correlation.

We then applied semipartial correlation (Pearson correlation) analysis to test if and to what extent the fMRI pattern dissimilarity structure could be explained by multiple predictor matrices reflecting the stimuli’s dissimilarity along several quantitative dimensions: numerosity, average item area, total field area, total surface area and density. Note that our design orthogonally manipulated numerosity, average item area and total field area. As a result, numerosity was partly correlated with density and total surface area (correlation between number and density predictor matrix = 0.84; between number and total surface area predictor matrix = 0.36). In order to control the potential impact of shared variance among our predictor variables and to ensure a more accurate assessment of each variable’s contribution, we employed semipartial correlation analysis (Abdi 2007). This statistical technique allowed us to disentangle the unique influence of each predictor variable from the common variance they share, thereby preventing an overestimation of their individual effects due to repetitive inclusion of shared components in our analysis. Thus, the coefficient of semipartial correlation between the neural RDM and the model RDM of interest represents the portion of unique variance shared between the neural RDM and the model RDM of interest while partialling out the effect of all other models. A schematic representation of this process is shown in Figure 2. To account for different levels of noise in different brain areas, we estimated the noise ceiling in all ROIs and normalized the semipartial correlation coefficients with the corresponding noise ceiling (Khaligh-Razavi et al., 2018; Al-Tahan & Mohsenzadeh, 2021). The noise ceiling was determined by correlating the mean across individual participant’s RDM and the group-averaged RDM (Nili et al., 2014).

**Figure 2.**
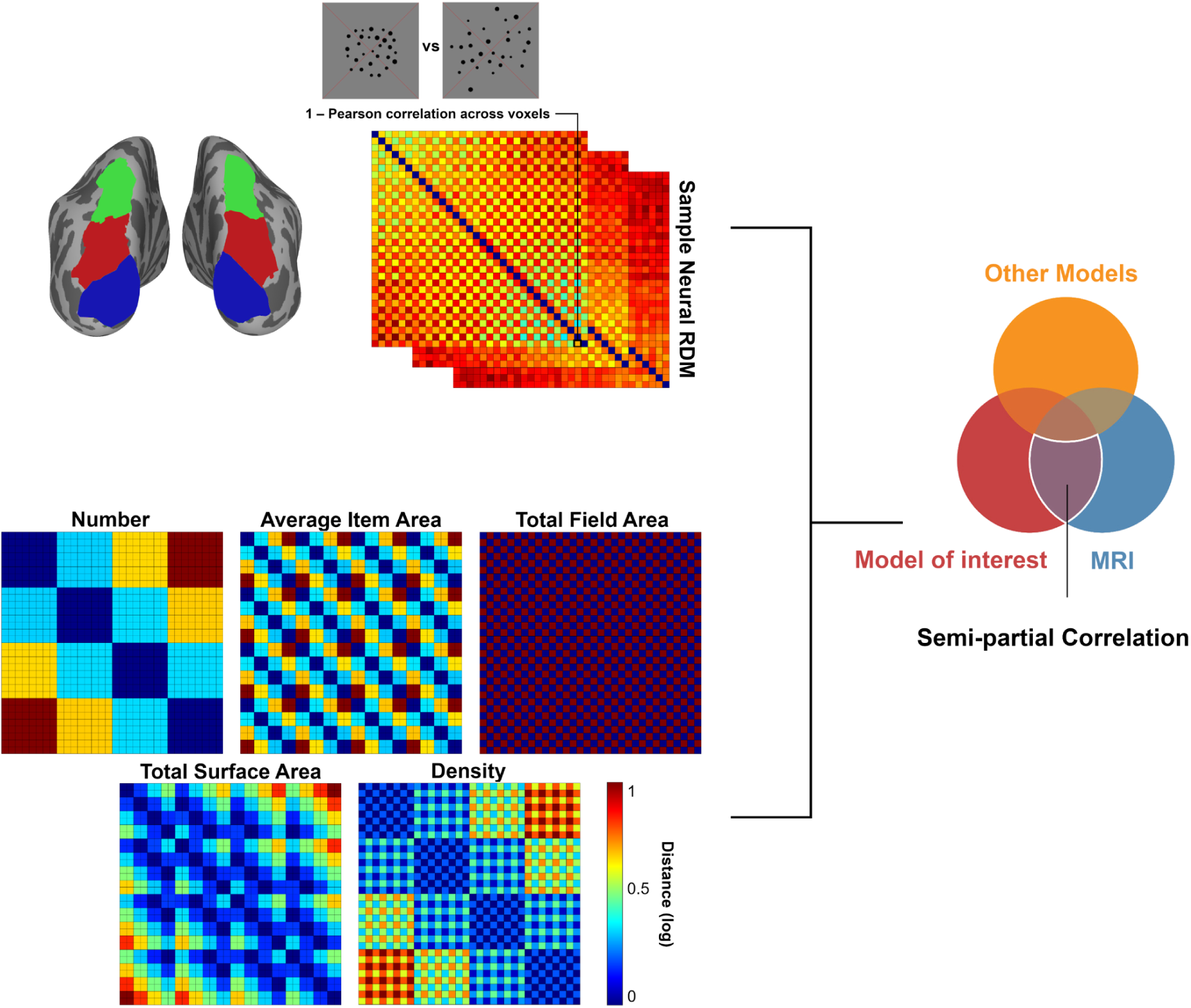
Neural representational dissimilarity matrices (RDM) derived from fMRI were entered into a semipartial correlation analysis. fMRI RDMs were created using 1 – Pearson correlation between the activations of voxels in that region for each pair of images. Five representational dissimilarity matrices, model RDMs, used as predictors in the semipartial correlation analysis. These matrices represent the logarithmic distance between pairs of stimuli in terms of number, average item area, total field area, total surface area, and density.

#### Surface-based ROI RSA

Following Castaldi et al. (2019), we selected several retinotopic regions of interest (ROIs) along the dorsal stream from a surface-based probabilistic atlas based on visual topography, averaging across the two hemispheres (Wang et al., 2015).

Moreover, because contrary to Castaldi et al. (2019) we recorded from the whole brain, we extended the analysis to the retinotopic regions of the ventral stream as defined by the same atlas (Wang et al., 2015). The ROIs in early visual areas were V1, V2 and V3. The ROIs along the dorsal stream were V3AB (merging V3A and V3B), V7, IPS12 (merging IPS1 and 2), IPS345 (merging IPS3, 4 and 5; Figure 3.A). These were further merged into two large ROIs that correspond to intermediate (V3A, V3B, and V7, also known as IPS0), and higher-level (IPS1 to IPS5) areas. The selected ROIs from the ventral stream were hV4, VO1, VO2, PHC1, PHC2 (Figure 3.B). The ventral ROIs were also further merged into two large intermediate (VO1 and VO2), and higher-level (PHC1 and PHC2) ROIs. In order to ease comparisons between ROIs we performed the subsequent analyses based on the 600 most active vertices (in the contrast “all sample stimuli > baseline”) in all ROIs (Mitchell et al., 2004). We choose the vertices from each individual ROI and large ROI separately.

**Figure 3.**
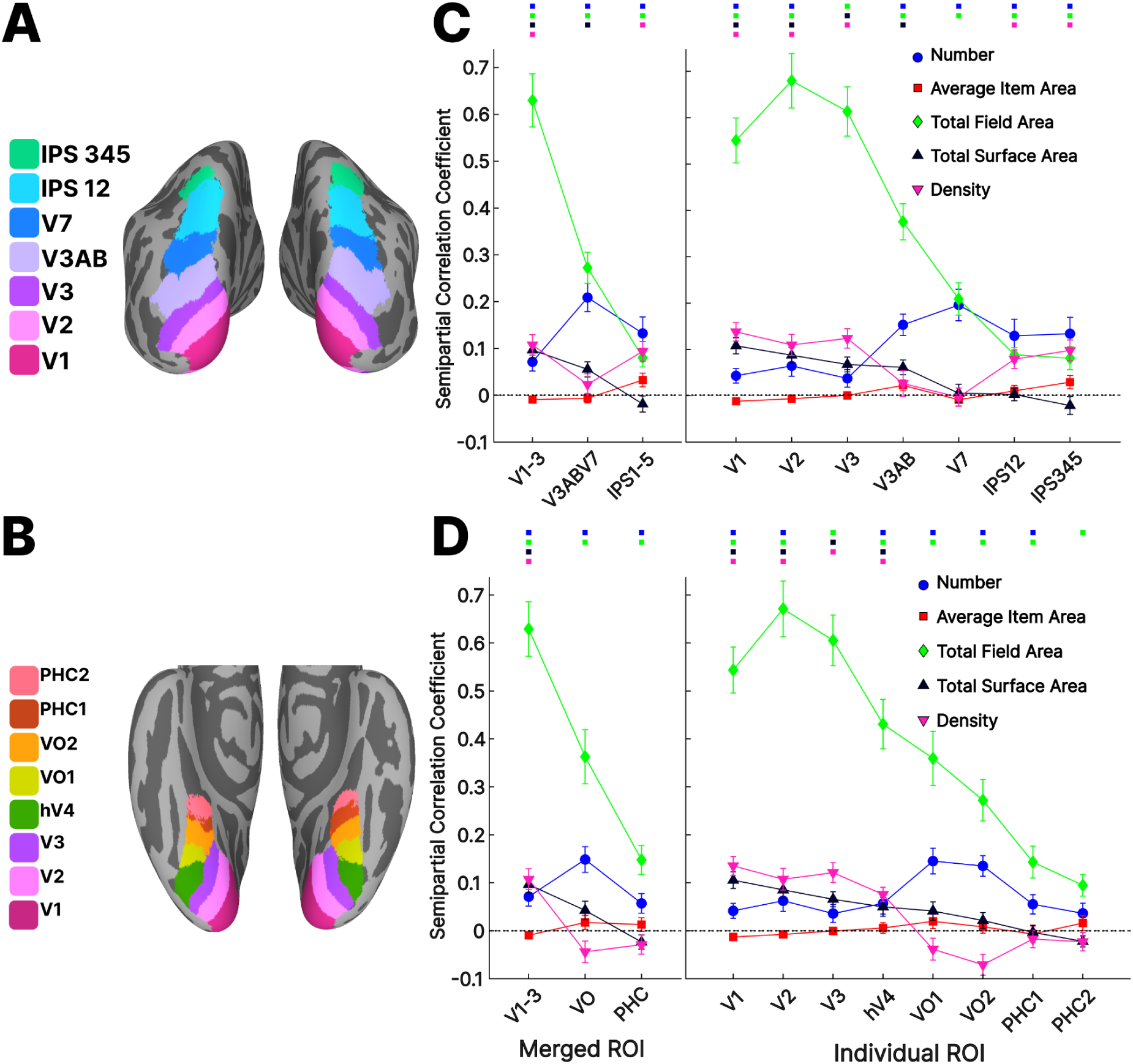
Color-coded ROIs for (A) dorsal and (B) ventral streams defined by the probabilistic atlas (Wang et al., 2015) are displayed on Freesurfer’s fsaverage inflated surface. (C) Semipartial correlation coefficients obtained from the representational similarity analysis for number, average item area, total field area, total surface area and density from predefined dorsal retinotopic ROIs. Number is represented along other visual features in almost all regions and is amplified in association areas (from V3AB to IPS). Data points show mean semipartial correlation coefficients across subjects (n = 31) ± standard error of the mean (SEM). The coloured points above the figure indicate where the effect significantly exceeds zero (p<0.01). (D) Semipartial correlation coefficients obtained from the representational similarity analysis for number and all other features from predefined retinotopic ventral ROIs after normalizing with the corresponding noise ceiling. Note that the results from early visual areas V1-3 are the same across panels as the ventral and dorsal components of these areas were merged into a single ROI. Within the ventral stream, number is amplified in intermediate areas (VO1 and VO2). Data points show mean semipartial correlation coefficients across subjects (n = 31) ± standard error of the mean (SEM). The coloured points above the figure indicate where the effect significantly exceeds zero (p<0.01).

The number of vertices (600) was chosen to be slightly lower than the maximum number of vertices across all ROIs (ranging from 661 to 1510). It is important to note, though, that when we selected all vertices across each ROI the results remained substantially unchanged. For the ROI RSA we used unsmoothed data but the results remained consistent when we smoothed the data. ROI-based RSA was implemented using the CoSMoMVPA MATLAB toolbox (Oosterhof et al., 2016) and custom-written code in MATLAB R2019. We used one-sample t-tests against zero across subjects to test the statistical significance of the Fisher-transformed semipartial correlation coefficients for each feature and ROI. We analyzed the effects of ROI and features with repeated measures analysis of variance (ANOVA).

#### Surface-based searchlight RSA

To find how numerical and non-numerical quantities are represented across the whole cortical surface, RSA was performed using a surface-based searchlight approach (Oosterhof et al., 2011), implemented using the CoSMoMVPA MATLAB toolbox (Oosterhof et al., 2016), the Surfing toolbox (Oosterhof et al., 2011), and a custom-written code in MATLAB R2019 (The MathWorks, Inc., Natick, MA). Each participant’s entire t-statistics brain map was smoothed with the same method we used to smooth the contrast map and underwent a searchlight (radius 6 mm along the cortical surface) procedure. A neural RDM was constructed using Pearson’s correlation. Similar to the ROI-based RSA, the semipartial correlation between the neural RDM and model RDMs were calculated and then mapped on the brain.

To identify vertices in which the Fisher-transformed semipartial correlation resulting from searchlight RSA was significantly above zero, a one-sample t-test was used across subjects. The result was then corrected using threshold-free cluster enhancement (TFCE; Smith & Nichols, 2009) using Monte Carlo simulations with 10,000 permutations implemented in the CoSMoMVPA MATLAB toolbox (Oosterhof et al., 2016). The resulting statistical map was thresholded at p < 0.01 (one-tailed) and was projected on the fsaverage surface for visualization.

#### Multidimensional scaling

The results from RSA show how each feature contributes to the variance in our data, in a hypothesis-driven manner. To further explore the latent structure of our neural data we complemented RSA with a data-driven approach, implementing multidimensional scaling (MDS; Kruskal, 1964) using the MATLAB function cmdscale. MDS spatially organizes the stimuli so that their relative distance mirrors the differences in the brain activity patterns they evoke. We used MDS and visualized the first two dimensions to investigate the neural representational geometry of our stimuli space both across and within ROIs. To compare the similarity of the neural representation of the stimulus space across ROIs, we vectorized the 32 x 32 group averages of individual neural RDMs reflecting the correlation across conditions within each of the 12 ROIs and then constructed a 12 × 12 RDM across ROIs. Then, to further explore the neural representational geometry within each stream, we applied the MDS on the group-average RDM across participants for each ROI. We also extended these analyses beyond the dorsal and ventral retinotopic regions including three additional clusters resulting from the whole brain searchlight map of regions encoding numerosity. To isolate these clusters we used group-constrained subject-specific (GCSS) analyses (Fedorenko et al., 2010; Julian et al., 2012), using custom-written MATLAB R2019 code (The MathWorks, Inc., Natick, MA) developed by Scott & Perrachione (2019; available at https://github.com/tlscott/make_parcels). The identification of these clusters involved a four-step process: Initially, for each participant, the Fisher-transformed semipartial correlation values resulting from the number searchlight maps were converted into z-scores. Subsequently, these z-scores were thresholded at p < 0.01 and binarized. Secondly, a probability map was generated by overlaying all binary maps. This resulting map was smoothed using a gaussian kernel of 6 mm FWHM, and vertices with contributions from fewer than ten subjects were set to zero. Thirdly, the watershed algorithm, as implemented in the SPM-SS toolbox (Nieto-Castañón & Fedorenko, 2012; https://www.nitrc.org/projects/spm_ss), was employed to locate local maxima. Clusters were defined around these local maxima and extended to neighboring vertices until a local minimum or a zero-valued vertex was encountered. The resulting parcels represent regions where multiple subjects exhibited suprathreshold activity, without the requirement that this activity occur in the exact same vertex across participants. Finally, the number of subjects contributing to each parcel was calculated, and parcels where more than 80% of subjects contributed were selected as the final parcels (Scott & Perrachione, 2019).

#### Code accessibility

All the code related to this study is available through GitHub: https://github.com/alireza-kr/numerosity_fmri-meg-cnn/

## Results

Thirty-one healthy adult volunteers were presented with visual arrays of dots orthogonally varying in numbers of items (6, 10, 17, 29), average item areas (0.04, 0.07, 0.12, 0.2 visual squares degree), and total field areas (9 or 13.5 visual degree diameter) while being scanned in a 3T MRI. Their task was to keep the number of dots in memory to compare it with an occasionally presented subsequent match stimulus. Behavioral performance on this numerosity comparison task was overall high (Mean=82.14%, SD=6.83%, Range=66.66% – 93.75%), indicating that subjects were attentive.

### A widespread network of cortical fronto-parietal regions overall involved in the task (univariate analyses)

We started the analysis of the functional imaging data by evaluating the overall regional activation during the experiment. Surface-based random-effects group analysis for sample stimulus against the implicit baseline revealed activation across both hemispheres in a wide set of regions extending both dorsally from early visual to parietal up to the postcentral gyrus and the precentral sulcus in the frontal cortex and ventrally including medial and lateral inferior occipito-temporal areas (Figure 1.C, thresholded at p < 0.01, TFCE corrected).

### Numerosity is encoded over and above the other features in retinotopic regions along both the dorsal and the ventral visual stream (ROI semipartial correlation RSA)

Following Castaldi et al. (2019), to disentangle the contribution of numerical and non-numerical features of the stimuli on the distributed patterns of activity of the BOLD signal, and to ask whether and where in the brain the representations of numerical and non-numerical features of the stimuli could be dissociated, we performed RSA (Kriegeskorte et al., 2008) using semipartial correlation, which ensures that the resulting coefficients reflect the unique variance explained by each model while partialling out the effect of all other models. We performed this analysis on 12 ROIs derived from a surface-based probabilistic atlas based on visual topography (Wang et al., 2015): 3 in early visual areas (V1, V2, V3), 4 along the dorsal stream (V3AB, V7, IPS12, IPS345) (replicating Castaldi et al., 2019), and 5 supplemental regions along the ventral stream (hV4, VO1, VO2, PHC1, PHC2).

The results indicate that the variance in brain activation patterns was significantly explained by number over and above all other non-numerical features in almost all regions (as shown by semi-partial correlations, where p<0.01 with the exception of V3 and PCH2) starting in early visual areas and reaching its highest explanatory power in higher-level regions V7-IPS along the dorsal stream (thus perfectly replicating Castaldi 2019) and in regions VO1-VO2 along the ventral stream.

Importantly, while number did account for unique variance, other predictors, such as total field area, exhibited stronger overall partial correlation coefficients with the neural signal in several regions, especially in early visual and in the ventral stream regions. An opposite pattern of results was seen for most non-numerical features that were maximally represented in early visual areas: total field area explained independent portions of variance in all regions, but maximally in V1-V3 and less so in higher-level regions of both streams; total surface area was significant only up to V3AB and hV4; density also was significant in V1-V3 and hV4, then ceased to explain a significant portion of the variance but regain some effect in IPS1-5. Average item area, instead, was never significant, neither in the dorsal nor in the ventral stream regions (Figure 3.C and 3.D).

To statistically evaluate the impact of the different features across the ROIs, we analyzed the semipartial correlation coefficients with four repeated measures ANOVAs (see below) with ROIs and features as factors. ROIs were considered both aggregated in three big regions, and separately for each of the individual regions, for the ventral and the dorsal stream separately. The significant two-way interaction between ROIs and features that we observed in all the 4 ANOVAs confirmed that the five features were differently encoded across ROIs across both the dorsal (for the three large regions: F(4.30,128.95)=36.680, p<0.001; for the individual seven regions: F(8.10, 243.08)=32.404, p<0.001) and the ventral stream hierarchy (for the three large regions: F(3.56, 106.91)=29.330, p<0.001; for the individual eight regions: F(9.56, 286.82)=27.513, p<0.001).

We then performed five one-way repeated measures ANOVAs on each feature across the ROIs. They revealed that, with the exception of average item area, all other stimuli features were encoded differently across ROIs along both streams (main effects of ROIs: p<0.01 for dorsal regions and p<0.01 for ventral regions, for both aggregated and individual ROIs)(average item area: three large ventral regions: F(1.91,57.40)=2.304, p=0.111; eight individual ventral regions: F(5.45,163.43)=1.362, p=0.238).

Taken together, these results indicate that numerosity is represented independently from other visual features along both dorsal and ventral retinotopic regions and that contrary to all other features it is amplified in associative areas (IPS and VO) in both streams. This is further supported by a positive linear regression of the semipartial correlation coefficient of number from V1 to IPS (β=0.02, p<1×10^−4^) and from V1 to VO (β=0.02, p<1×10^−4^), with a negative linear regression coefficient observed across regions for all other features from V1 to IPS (TFA: β=-0.09, TSA: β=-0.02, Density: β=-0.01, p<0.01) and from V1 to VO (TFA: β=-0.06, TSA: β=-0.02, Density: β=-0.04, p<0.001), except average item area (p<0.05).

### Numerosity is encoded beyond retinotopic regions in both the dorsal and the ventral visual stream (Searchlight semipartial correlation RSA)

After having replicated and extended the results of Castaldi et al. (2019) to the ventral visual retinotopic regions, we broadened our analyses across the entire cortical surface through a surface-based searchlight RSA. Results revealed that numerosity explained independent variance in several regions spread across the cortical surface (Figure 4), extending well beyond the previously defined retinotopic ROIs. These regions include the mid and anterior parietal cortex, both superior, inferior, and anterior to the intraparietal sulcus, the parieto-occipital cortex, the precentral gyrus and superior frontal sulcus. Along the ventral stream, these regions extended anteriorly and laterally in the mid inferior temporal lobe compared to the retinotopic regions VO1 and VO2. Relating our results to previously described functional parcellations of the ventral stream, our large ventral numerosity-related region [RH: 38, −69, −14; LH: −46, −69, −1] appears to potentially overlap with the ‘number form area’ reported by Yeo et al. (2017), located at MNI coordinates [RH: 55, −50, −12], and may also potentially extend further and include the “visual word form area-1”, the “fusiform face area” and the “fusiform body area”, as defined by the functional atlas by Rosenke and colleagues (Rosenke et al., 2020). Thus, numerosity is also encoded, irrespective of other visual features, in non topographically organized areas in the parietal, occipito-temporal, as well as in the frontal lobes. On the contrary, all non-numerical features (apart from average item area, for which we could not find any regions representing it) explained variance mainly in the early visual cortex (Figure 4).

**Figure 4.**
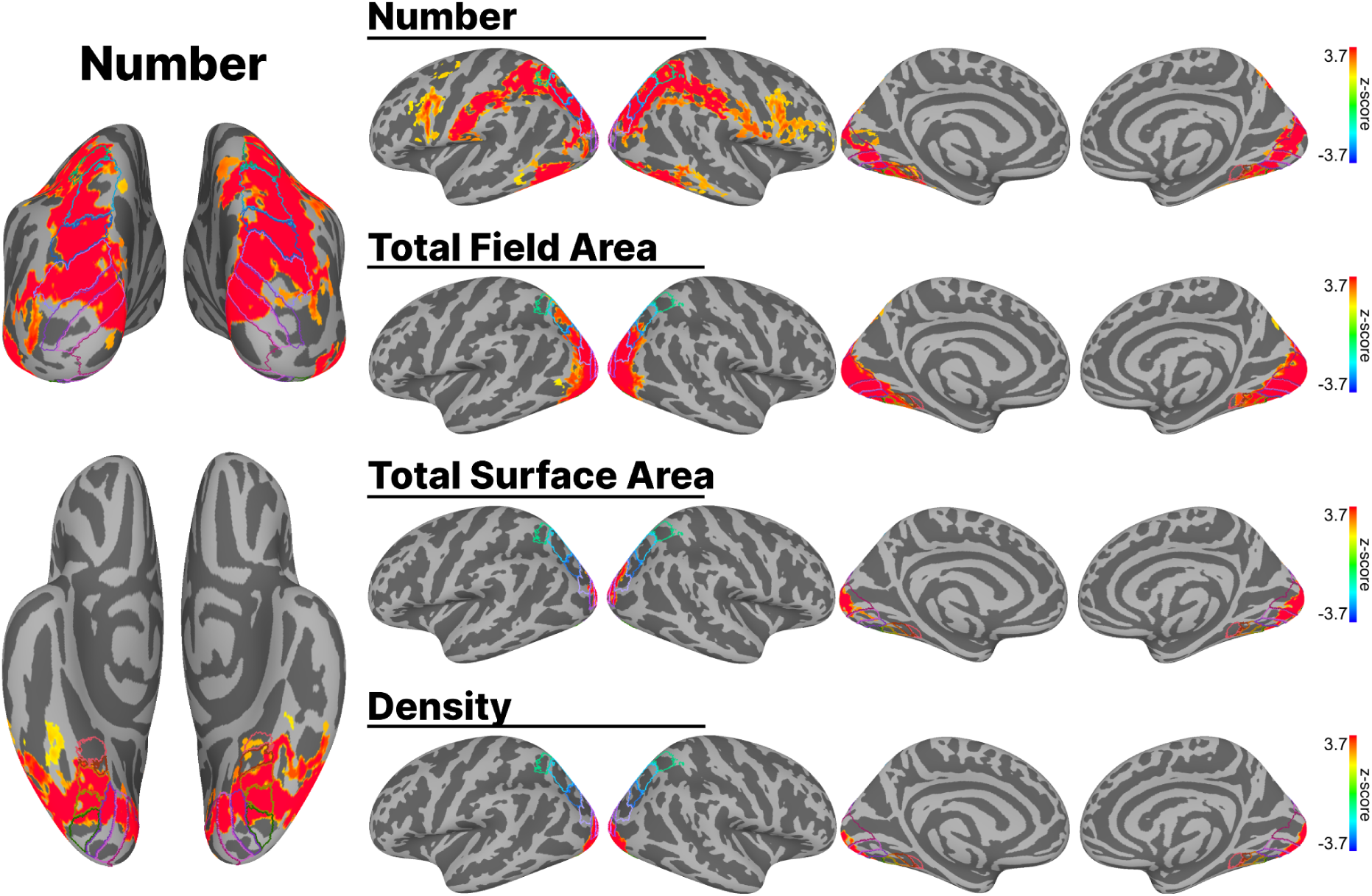
Surface-based searchlight representational similarity analysis searchlight (RSA) results obtained from the surface-based group analysis (n = 31). The maps show how patterns of activity across the cortical surface are captured by each model of interest (Number, Total Field Area, Total Surface Area, and Density) while partialling out the effect of other models. Activation maps are thresholded at p < 0.01, TFCE corrected, and displayed on Freesurfer’s fsaverage surface with colored outlines identifying ROIs along the early visual (V1, V2, V3), the dorsal (V3AB, V7, IPS12, IPS345), and the ventral stream (hV4, VO1, VO2, PHC1, PHC2) from a surface-based probabilistic atlas (Wang et al., 2015).

#### The neural representational geometry of numerosity differs between early visual and association cortices (Multidimensional Scaling)

To further explore whether the neural representational geometry of our stimulus space was similar or differed across early and associative regions and across associative regions of the two streams, we used multidimensional scaling (MDS), a data-driven approach that recovers a low-dimensional representation of the neural similarity structure and that can potentially reveal unpredicted coding properties. We applied MDS both across and within ROIs.

The MDS across all retinotopic ROIs (Figure 5.A) revealed three clusters: one grouping together all early visual regions, the other all ventral regions, and the third one all the dorsal regions, suggesting that these three groups of regions represent the stimulus space differently. This result prompted us to further investigate the neural representational geometry of each ROI separately (see Figure 5.B). The reconstructed neural geometry of the early visual areas indicated a clear ordered representation of numerosity along both the first 2 dimensions of the stimulus space, akin to a number line, as well as a clear separation between stimuli with large and small total field area, coherently with the RSA results. While this pattern remained approximately constant along the ventral stream ROIs up to VO2, it changed as we approached the higher ROIs along the dorsal stream (IPS12 and IPS345): here the separation based on total field area decreased and a curved pattern around the midpoints of the number continuum emerged. This parietal curved pattern (a bent “number line”, where along the second dimension central numbers are encoded separately from extreme numbers) was not at all evident in the ventral stream retinotopic association areas VO1 and VO2, even if numerosity was equally well represented across streams.

**Figure 5.**
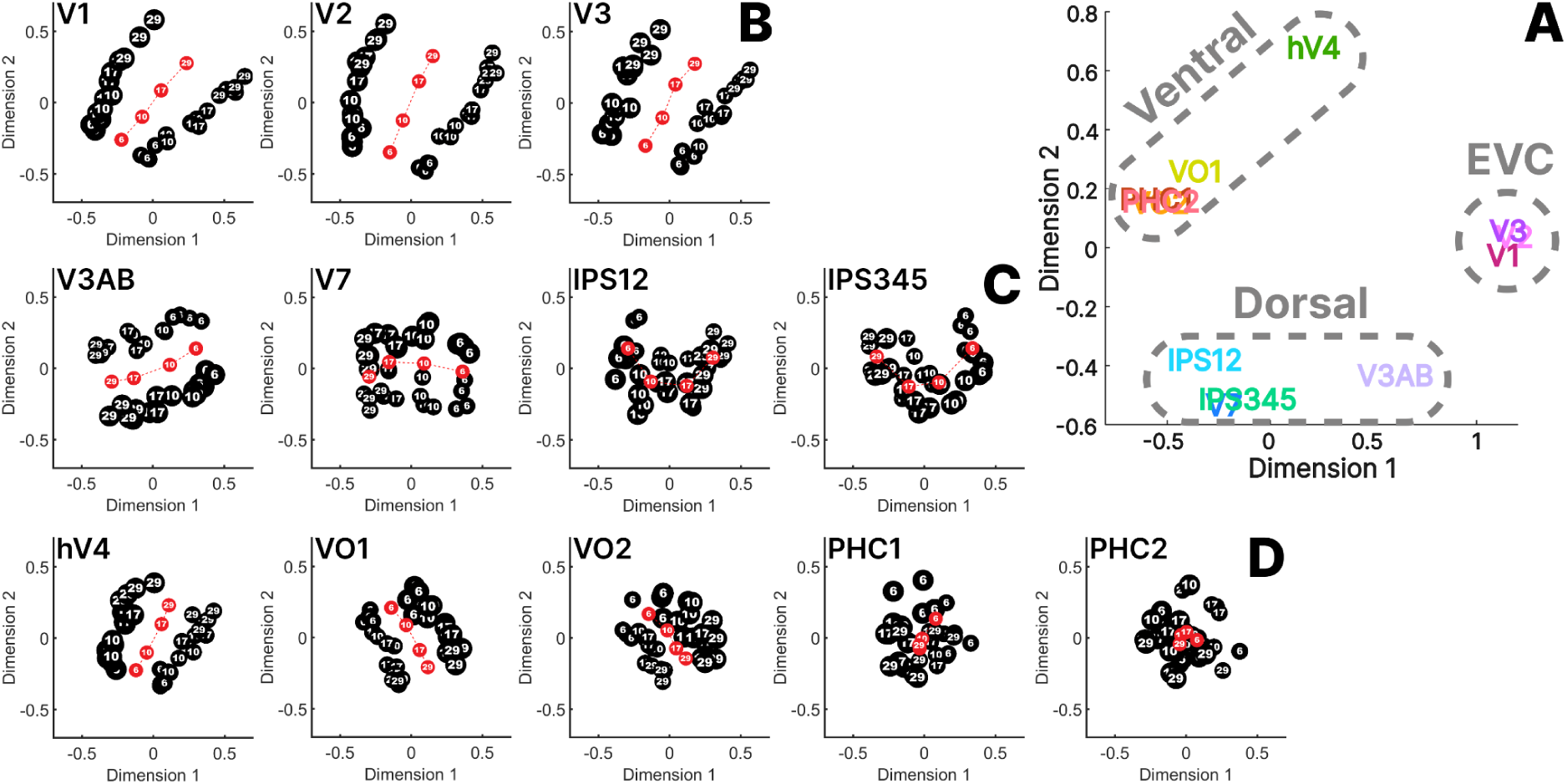
(A) Multidimensional scaling (MDS) reveals similarity of the representational structures between the predefined ROIs in a two-dimensional space. Here the proximity between any two ROIs indicates how similar their representation of the stimuli is. Three clusters of regions become apparent. MDS reveals representational similarities between stimuli in a two-dimensional space for regions in the (B) EVC, (C) dorsal regions, and (D) ventral regions. The black circles represent the 32 stimuli. They are labeled according to their numerosity (6, 10, 17, 29), and scaled in size to represent stimuli with small total field area (small circles) and larger total field area (large circles). The red circles indicate the average coordinates of each number.

Interestingly, however, this difference in the neural manifold across the dorsal and the ventral streams was reduced when we focussed on regions outside the retinotopic ROIs. Within the group-level maps resulting from the searchlight analysis we isolated the most reproducible parcels across subjects (group-constrained subject-specific (GCSS) analyses, see Methods) that resulted in three clusters: two parietal (one anterior and one posterior, labeled “NPC1” (as in Numerosity Parietal Cortex) and NPC2, and one occipito-temporal, labeled “NTO” (as in Numerosity Temporo-Occipital). We adopted these naming conventions following the one used by Harvey & Dumoulin (2017) because of the apparent overlap between our clusters and the ones reported by those authors. For a direct comparison with the Harvey and Dumoulin numerotopic maps, the MNI x,y,z peak coordinates of our ventral parcels were [RH: 38, − 69, − 14; LH: − 46, −69, − 1], while the centers of NTO as found by Harvey and Dumulin were [RH: 44(7), − 75(1), − 4(3); LH: −42(3), −77(3), −3(8)]. Our parietal peaks of cluster NPC1 were [RH: 35, − 54, 56; LH: − 39, −41, 43] while those from Harvey and Dumulin were [RH: 22(5), −61(7), 60(5); LH −22(4),−59(11), 61(8)]; our peaks for NPC2 were [RH: 46, − 25, 40; LH: − 63, − 23, 28] while those from Harvey and Dumulin were [RH 33(3), − 40(4), 52(7), LH − 38(3), − 43(8), 48(8)]. These comparisons have to be taken cautiously, however, as in both Harvey and Dumoulin and the present research the clusters were very large and not necessarily well represented by the localization of the peak voxel.

We then completed our analyses by performing MDS on the activity in those clusters and the results revealed a clear curved number line structure both in the parietal regions NPC1 and NPC2 as well as in the inferotemporal region NTO. These results suggest that while all along the dorsal stream the representational geometry of numbers looks like a bended line, along the ventral stream there is a mixture of representational geometries across regions, taking the form of straight or curved number lines. In order to assess whether additional variance or hidden structures could be captured from additional latent dimensions of the MDS, we also investigated the third dimension of the MDS; as detailed in the Supplementary Materials (Section S1), however, we found that the third dimension did not explain significant more variance, nor it did reveal additional features of the neural representational geometry that was not already captured by the first two dimensions. In sum, while in early visual cortex regions and in most retinotopic regions of the ventral stream the neural manifold was essentially a straight number line, in all associative regions along the dorsal stream and in one lateral patch along the ventral stream the manifold consisted in a curved structure where in one dimension numerosities are well ordered while in the other dimension the extreme ones are encoded separately from intermediate ones (Figure 6).

**Figure 6.**
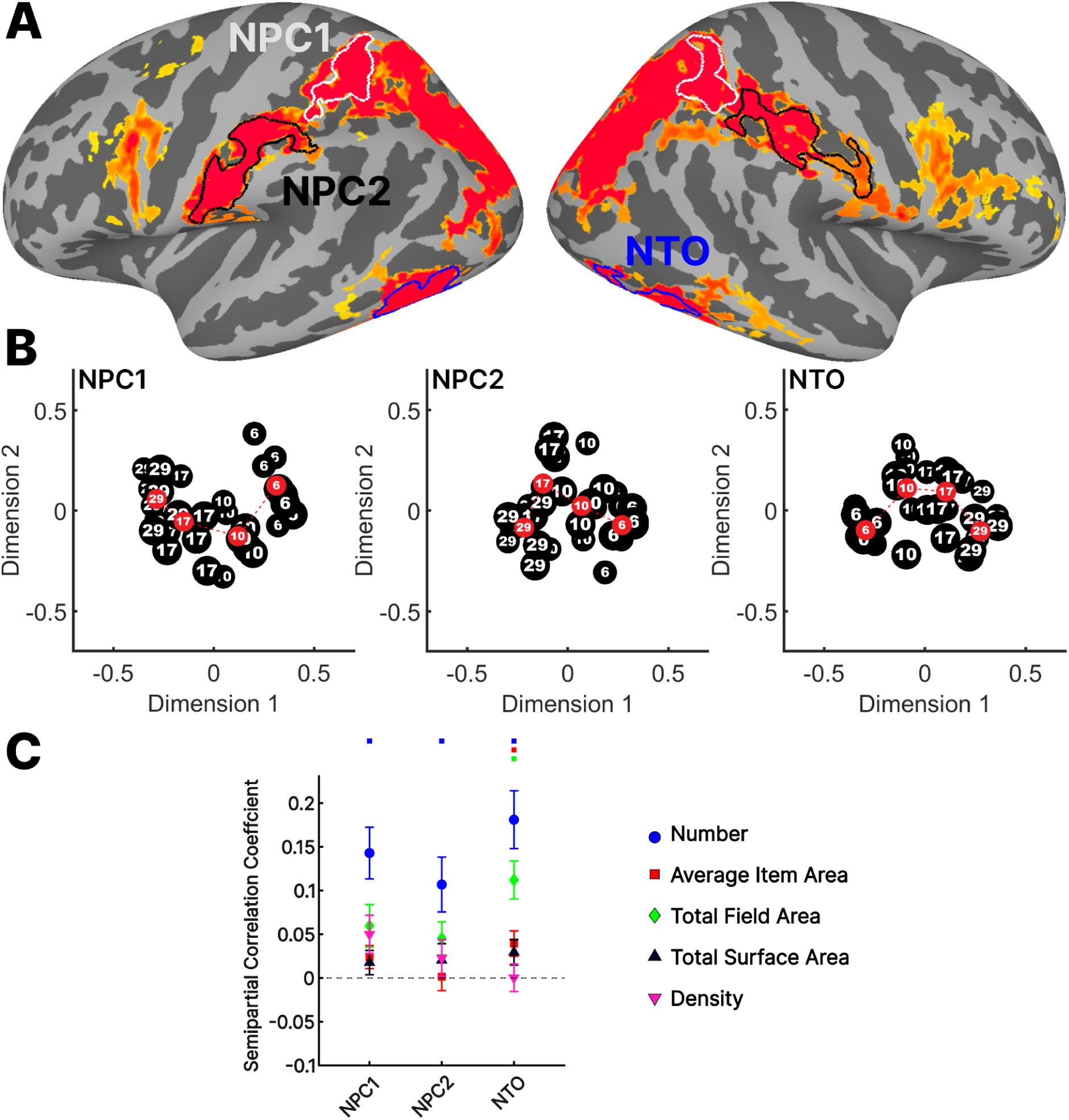
(A) ROIs (NTO, NPC1, NPC2) chosen using group-constrained subject-specific (GCSS) analysis from outside the retinotopic regions of interest, as defined in the probabilistic atlas by Wang et al. (2015), were visualized on the number activation map derived from the searchlight analysis. (B) Multidimensional scaling (MDS) reveals representational similarities between stimuli in a two-dimensional space for regions in the dorsal (NPC1, and NPC2) and the ventral (NTO) stream. (C) Semipartial correlation coefficients obtained from the representational similarity analysis for number and all other features from regions in the dorsal (NPC1, and NPC2) and the ventral (NTO) stream

## Discussion

We used Representational Similarity Analysis (RSA) and Multidimensional Scaling (MDS) to investigate whether, where and how the human adult brain represents numerosity independently from other visual features using fMRI and a numerosity estimation task.

The region of interest and searchlight RSA showed that numerosity and the other visual properties of sets are represented independently from one-another in several brain regions, consistent with the existence of partially independent channels for different quantitative properties of visual sets. However, while the representations of the non-numerical visual quantitative features such as total field area, total surface area and density tended to remain confined to early visual areas (with the notable exception of average item area, which we did not find to be encoded in any region), the representation of numerosity was present more widely across the cortex, far exceeding the retinotopic regions, and amplified in association areas not only along the dorsal stream, as often previously reported in the literature, but also along the ventral visual stream of both hemispheres. Starting from early visual regions (V1-V3), numerosity is progressively amplified both within and outside retinotopic regions along both streams, extending to both anterior, superior, and inferior to the IPS, and both anterior and lateral to the occipito-temporal areas VO. While most previous studies investigating large numerosity representations reported activation around the parietal cortex, there is extremely little evidence for large numerosity representation along the ventral stream. Notably, Cai et al. (2021), using the population receptive field (pRF) mapping method (Harvey & Dumoulin, 2017), also found large numerosity representations (in the form of numerotopic maps) in association regions of both the dorsal and ventral streams. However, this study had two important limitations. First, in their stimuli, object size and density were correlated with number, thus we cannot exclude that their results reflected an ordered representation of size instead of number. Second, they presented stimuli in a strictly ordered manner (with numerosity either increasing or decreasing), a design that can introduce expectations and attentional biases whose effects cannot be easily separated from the sensory representation of the stimuli themselves. Indeed, expectations about specific categories can lead to an increase in BOLD signals within category-selective regions, even in the absence of stimuli. Set aside the study by Cai, the fact that the numerosity representation along the ventral stream has been largely neglected in the previous neuroimaging literature could be explained by the fact that, due to prior biases towards parietal cortex, some only scanned a limited brain volume centered on the dorsal visual pathway (e.g., Lasne et al., 2019; Castaldi et al., 2019), while those recording the whole brain often restricted their analyses on parietal cortex regions (e.g., Bulthè et al., 2015); others, finally, may have been limited in their power to detect such ventral activation due to the use of passive or minimally demanding tasks (e.g., Piazza et al., 2004). However, also in light of the neuropsychological literature that consistently associates deficits in numerosity processing with parietal rather than occipito-temporal cortex damage (Warrington & James, 1967; Lemer et al., 2003), our report of a robust response to numerosity in occipito-temporal regions pave the way for future studies investigating its role in numerosity perception. We propose two potential hypotheses based on this result, which can be tested in future research. The first one sees the representation of numerosity in the ventral stream as the result of formal math education and of the functional connectivity between this region and the parietal regions (Kay & Yeatman, 2017; Conrad et al., 2022). This view stems from two observations: 1) young children are endowed with a precocious sense of numerosity that seems to mainly emerge from a parietal cortex circuit (Cantlon et al., 2006; Izard et al., 2008; Hyde et al., 2010), 2) during math education a specific portion of the ventral visual stream functionally specializes in the recognition of digits, giving rise to the Number Form Area (NFA; Amalric & Dehaene, 2016; Grotheer et al. 2018; Kersey et al., 2019; Kutter et al., 2022; see for a review: Yeo et al., 2017). Also, the cortical areas corresponding to the NFA appear as potentially partially overlapping with those we find to represent numerosity along the ventral stream. However, further work at the individual level is necessary to delineate the specific contributions of these areas.

According to this view, when a cortical patch in the ventral stream develops its response to number symbols, it concurrently increases its connectivity with the parietal regions encoding numerosity. This increased connectivity might cause numerosity representations to be broadcasted between parietal cortex and the NFA, such that both regions eventually come to represent both arabic digits and the associated numerosities (Piazza et al., 2007; Eger et al., 2009). Interestingly, and in line with this idea, a recent study performed on a few subjects and using the pRF approach indicated that neuronal populations tuned to small numerosity in the ventral occipitotemporal cortex region NTO also respond to arabic digits (Cai et al., 2023). This hypothesis predicts that the occipito-temporal response to numerosities may not be present in preschoolers, and that it develops as a result of formal math education (Dehaene-Lambertz et al., 2018; Kersey et al., 2019). However, this hypothesis must be considered as highly tentative as the precise anatomical location of the NFA region remains rather poorly determined, and the overlap between our NTO map encoding numerosity and the NFA as defined in a recent meta-analysis (Yeo et al., 2017) is only partial. Moreover, we do not know whether our ventral numerosity-responsive regions NTO would also encode digits, as we did not expose subjects to such stimuli.

Another possible interpretation, not necessarily exclusive with the first one, posits that both the dorsal and the ventral stream possess sufficient computational resources to independently represent numerosity, and that different representations of numerosity might serve different functional purposes. This view would also be in line with some results from the literature on visuo-spatial attention, where, in both monkeys and humans, it was shown that attentional control networks, containing attentional priority location-based maps of the stimuli, and typically associated with a dorsal fronto-parietal circuit, also include a ventro-temporal cortical node, the phPTG, whose location seems compatible with that of our ventral activation patch (Stemmann & Freiwald, 2019; Sani et al., 2021). Prior research on numerosity representation had indeed speculated that it could emerge from a saliency-map type of system, where attention is distributed on as many locations as there are items in the display, providing a good proxy for approximate numerosity representation (e.g. Knops et al., 2014; Verma & Sengupta, 2023). In contrast with the first hypothesis, this hypothesis predicts that numerosity is represented in both streams independent from and prior to math education.

To investigate the differences in how various numerosity-coding regions represent numerosity we applied Multidimensional Scaling, a hypothesis-free dimensionality reduction approach that can uncover latent patterns in the neural data that may otherwise remain invisible with model-based approaches. In early visual cortices (V1-V3), we observed a clear linear rank-ordering of numerosities, which was consistently present in all dimensions tested. In contrast, associative regions along both the dorsal and the ventral stream, displayed a curved manifold, with two main axes: one ordering the numerosities and the other(s) separating extreme from middle values (see Supplementary Material S1 and S2). The fact that associative and early visual regions are characterized by two different neural representational geometries could be explained in two ways.

The first one is that while early visual regions solely encode the quantitative aspect of the stimuli (numerical magnitude), associative regions represent both numerical magnitude (encoded in the first dimension of the MDS of the neural RDM) and at the same time the relative status of each number within the rank, encoding extreme values differently from the intermediate ones (corresponding to the second dimension of the MDS of the neural RDM). Such differential encoding of extreme vs. middle values would nicely align with prior similar findings that during rank-ordering tasks (Nelli et al., 2023), the posterior parietal cortex encodes the stimuli with a bended “number line”. This would suggest that magnitude and order are represented through a similar neural schema, fitting with the proposal that the posterior parietal cortex abstracts relational information and compresses decision-relevant variables into low-dimensional representations (Summerfield et al., 2020). However, we must acknowledge a major difference between our experiment and the one by Nelli et al. (2023), in that in our experiment we analysed the brain activity evoked solely during the encoding of the stimuli, prior to and independent from the comparative judgement, thus devoid from explicit decisions. Indeed, in our experiment subjects were primarily passively encoding numerosity, with only a very small subset of trials in which they were asked to perform a one-back number comparison task (which we later excluded from the analysis). It is possible that even if subjects were not required to perform an explicit decision during the fMRI recording they nevertheless activated a response classifying the sets as having extreme (smallest/largest), or intermediate numerosities. This would be coherent with previous reports that associative, but not primary areas host numerosity representations that are explicitly read out for numerical decisions making (Lasne et al., 2019; Moscoso et al., 2021). For example, Lasne et al. (2019) showed adult subjects visual set of different number of dots and observed that while numerosity could be equally well decoded by the BOLD activity in both early visual and parietal cortex, the inter-individual variability in numerosity comparison performance was predicted by the decoding accuracy of the BOLD signal in parietal, but not in early visual cortex. Currently, however, no studies directly compared the behavioral relevance of the parietal and the occipito-temporal representations of numerosity, thus this remains a question that future studies should resolve.

Together with this interpretation of the curved vs. linear structure of the neural representations of number across brain regions, we must acknowledge an alternative one, that more directly relates to the tuning scheme of the underlying neurons: according to this interpretation, the linear structure observed in the MDS of the early visual regions reflects a monotonic neural code for number, while the curved structure observed in the MDS of parietal regions reflects numerosity-tuned responses. This interpretation would align with findings from Paul et al. (2022), who, using a receptive field mapping approach, found that while the BOLD response to numerosity in early visual areas, particularly V1, follow a monotonic pattern— a consistent increase or decrease with numerosity without peaking at a specific number— higher-order regions, such as the lateral occipital and parietal cortices, exhibit numerosity-tuned responses, where different voxels peak at specific numerosities with a gaussian-like response function (activity peaking at a preferred numerosity and decreasing for both smaller and larger values). Could this difference account for our different representational geometries as extracted through multivoxel pattern similarity approach? While the relationship between tuning functions and representational geometry is complex — since differently tuned units can induce similar representational geometries at the population level (Kriegeskorte & Wei, 2021; Khosla & Williams, 2023) — we conducted some simple simulations. In those simulations, we analysed the activity of a population of synthetic voxels, hypothesizing that each one would respond preferentially to one numerosity with either a monotonic or tuned response. Initially we assumed that the same proportion of voxels would code each numerosity and then we varied the number of voxels tuned to each number range based on biologically plausible results (Cai et al., 2021), noticing that the results did not vary. We performed RSA on those modelled responses and used MDS to visualize the resulting representational geometries. As detailed in the Supplementary Materials (Section 2), our simulations show that, under certain parameter settings (e.g., relatively small tuning widths not constrained by biological plausibility), monotonic responses yield a linear arrangement, whereas numerosity-tuned responses give rise to a curved structure in MDS. To be more biologically plausible, we also ran the simulation with a larger standard deviation—following the estimates of Cai et al. (2021)—but these results deviated from those obtained with the smaller SD and failed to produce the curved MDS configuration. The cause of this discrepancy remains unclear. Although grounded in oversimplified assumptions and parameter choices that are not always biologically plausible, our findings support the notion that a curved “number line” in fMRI data could emerge from numerosity-tuned neural populations, while a linear representation may reflect aggregated monotonic responses. However, given these model simplifications and sensitivity to parameter choice, this interpretation remains provisional and warrants further investigation.

In summary, this study demonstrates that numerosity representations, independent of other non-numerical visual features, are broadly present in the brain starting from early visual areas and further amplified following a gradient from early visual areas, reaching its peak in association cortices along both the dorsal (up to anterior parietal cortex) and ventral streams (up to lateral and inferior occipito-temporal cortex), challenging the mainstream view of a preferential role of the dorsal stream in numerosity representation (Dehaene and Cohen, 1995; Dehaene et al., 2003).

Model-free dimensionality reduction analyses further indicated that the neural representational geometry of numerosity differs substantially across early and association regions: the former encode numerosity solely according to numerical rank order while the latter, especially along the dorsal stream, are characterized by a curved manifold. These findings are compatible with two not necessarily exclusive interpretations: early visual regions encode solely number while association areas encode both number and their status on a mental line (extreme vs. intermediate values), or early visual regions code for number with a monotonic code while associative regions with a numerosity-tuned code. Future work will disentangle between the two.

## Author contribution

AK: Conceptualization, Data Curation, Formal Analysis, Investigation, Methodology, Software, Visualization, Writing – Original Draft Preparation, Writing – Review & Editing;

MP: Conceptualization, Supervision, Validation, Writing – Original Draft Preparation, Writing – Review & Editing, Project Administration, Funding Acquisition;

EC: Conceptualization, Supervision, Writing – Review & Editing;

EE: Conceptualization, Supervision, Writing – Review & Editing;

## Acknowledgments

MP and AK acknowledge the Italian ministry for Education, University and Research (MIUR) for the Departments of Excellence Grant supporting a PhD scholarship to AK and the Center for Mind and Brain Sciences (CIMeC) for supporting the research. For this work AK was awarded the 42° European Workshop on Cognitive Neuropsychology (https://www.ewcn.eu/) prize. MP and EC were funded by the Italian Ministry of Education, University, and Research under the PRIN2022 program (MP: grant number 2022EBC78W - ‘Sense of number vs. sense of quantity: modeling, neuroimaging, behavior’; EC: grant number 2022CCPJ3J - ‘RIGHTSTRESS - Tuning arousal for optimal perception’).

## References

Abdi, H. (2007). Part and partial correlations. In N. J. Salkind (Ed.), Encyclopedia of Measurement and Statistics (pp. 736–740).

Al-Tahan, H., & Mohsenzadeh, Y. (2021). Reconstructing feedback representations in the ventral visual pathway with a generative adversarial autoencoder. PLOS Computational Biology, 17(3), e1008775. 10.1371/journal.pcbi.1008775

Amalric, M., & Dehaene, S. (2016). Origins of the brain networks for advanced mathematics in expert mathematicians. Proceedings of the National Academy of Sciences of the United States of America, 113(18), 4909–4917. 10.1073/pnas.1603205113

Anobile, G., Cicchini, G. M., & Burr, D. (2013). Separate mechanisms for perception of numerosity and density. Psychological Science, 25(1), 265–270. 10.1177/0956797613501520

Ansari, D., & Dhital, B. (2006). Age-related Changes in the Activation of the Intraparietal Sulcus during Nonsymbolic Magnitude Processing: An Event-related Functional Magnetic Resonance Imaging Study. Journal of Cognitive Neuroscience, 18(11), 1820–1828. 10.1162/jocn.2006.18.11.1820

Behzadi, Y., Restom, K., Liau, J., & Liu, T. T. (2007). A component based noise correction method (CompCor) for BOLD and perfusion based fMRI. NeuroImage, 37(1), 90–101. 10.1016/j.neuroimage.2007.04.042

Brainard, D. H. (1997). The Psychophysics Toolbox. Spatial Vision, 10(4), 433–436. 10.1163/156856897x00357

Bulthé, J., de Smedt, B., & op de Beeck, H. (2014). Format-dependent representations of symbolic and non-symbolic numbers in the human cortex as revealed by multi-voxel pattern analyses. NeuroImage, 87, 311–322. 10.1016/j.neuroimage.2013.10.049

Bulthé, J., de Smedt, B., & op de Beeck, H. P. (2015). Visual Number Beats Abstract Numerical Magnitude: Format-dependent Representation of Arabic Digits and Dot Patterns in Human Parietal Cortex. Journal of Cognitive Neuroscience, 27(7), 1376–1387. 10.1162/jocn_a_00787

Burr, D., & Ross, J. (2008). A Visual Sense of Number. Current Biology, 18(6), 425–428. 10.1016/j.cub.2008.02.052

Burr, D., Anobile, G., & Arrighi, R. (2018). Psychophysical evidence for the number sense. Philosophical Transactions of the Royal Society B, 373(1740), 20170045. 10.1098/rstb.2017.0045

Cai, Y., Hofstetter, S., Van Dijk, J. A., Zuiderbaan, W., Van Der Zwaag, W., Harvey, B. M., & Dumoulin, S. O. (2021). Topographic numerosity maps cover subitizing and estimation ranges. Nature Communications, 12(1). 10.1038/s41467-021-23785-7

Cai, Y., Hofstetter, S., & Dumoulin, S. O. (2023). Nonsymbolic numerosity maps at the occipitotemporal cortex respond to symbolic numbers. The Journal of Neuroscience, 43(16), 2950–2959. 10.1523/jneurosci.0687-22.2023

Cantlon, J. F., Brannon, E. M., Carter, E., & Pelphrey, K. A. (2006). Functional imaging of numerical processing in adults and 4-y-Old children. PLOS Biology, 4(5), e125. 10.1371/journal.pbio.0040125

Castaldi, E., Mirassou, A., Dehaene, S., Piazza, M., & Eger, E. (2018). Asymmetrical interference between number and item size perception provides evidence for a domain specific impairment in dyscalculia. PLOS ONE, 13(12), e0209256. 10.1371/journal.pone.0209256

Castaldi, E., Piazza, M., Dehaene, S., Vignaud, A., & Eger, E. (2019). Attentional amplification of neural codes for number independent of other quantities along the dorsal visual stream. ELife, 8. 10.7554/elife.45160

Castaldi E., Vignaud A., & Eger E. (2020). Mapping subcomponents of numerical cognition in relation to functional and anatomical landmarks of human parietal cortex. Neuroimage. 221:117210. 10.1016/j.neuroimage.2020.117210

Castelli, F., Glaser, D. E., & Butterworth, B. (2006). Discrete and analogue quantity processing in the parietal lobe: A functional MRI study. Proceedings of the National Academy of Sciences of the United States of America, 103(12), 4693–4698. 10.1073/pnas.0600444103

Chung, M. K., Robbins, S., Dalton, K. M., Davidson, R. J., Alexander, A. L., & Evans, A. C. (2005). Cortical thickness analysis in autism with heat kernel smoothing. NeuroImage, 25(4), 1256–1265. 10.1016/j.neuroimage.2004.12.052

Conrad, B., Pollack, C., Yeo, D. J., & Price, G. R. (2022). Structural and functional connectivity of the inferior temporal numeral area. Cerebral Cortex, 33(10), 6152–6170. 10.1093/cercor/bhac492

Cox, R. W. (1996). AFNI: Software for analysis and Visualization of Functional Magnetic Resonance neuroimages. Computers and Biomedical Research, 29(3), 162–173. 10.1006/cbmr.1996.0014

Decarli, G., Zingaro, D., Surian, L., & Piazza, M. (2023). Number sense at 12 months predicts 4-year-olds’ maths skills. Developmental Science, 26(6). 10.1111/desc.13386

Dehaene-Lambertz, G., Monzalvo, K., & Dehaene, S. (2018). The emergence of the visual word form: Longitudinal evolution of category-specific ventral visual areas during reading acquisition. PLOS Biology, 16(3), e2004103. 10.1371/journal.pbio.2004103

Dehaene, S., & Cohen, L. (1995). Towards an anatomical and functional model of number processing. Mathematical Cognition, 1, 83–120.

Dehaene, S., Piazza, M., Pinel, P., & Cohen, L. (2003). Three parietal circuits for number processing. Cognitive Neuropsychology, 20(3–6), 487–506. 10.1080/02643290244000239

DeWind, N. K., Park, J., Woldorff, M. G., & Brannon, E. M. (2019). Numerical encoding in early visual cortex. Cortex, 114, 76–89. 10.1016/j.cortex.2018.03.027

Ditz, H. M., & Nieder, A. (2016). Numerosity representations in crows obey the Weber–Fechner law. Proceedings of the Royal Society B: Biological Sciences, 283(1827), 20160083. 10.1098/rspb.2016.0083

Eger, E., Michel, V., Thirion, B., Amadon, A., Dehaene, S., & Kleinschmidt, A. (2009). Deciphering Cortical Number Coding from Human Brain Activity Patterns. Current Biology, 19(19), 1608–1615. 10.1016/j.cub.2009.08.047

Eger, E. (2016). Neuronal foundations of human numerical representations. Progress in Brain Research, 1–27. 10.1016/bs.pbr.2016.04.015

Faye, A., Jacquin-Courtois, S., Reynaud, E., Lesourd, M., Besnard, J., & Osiurak, F. (2019). Numerical cognition: A meta-analysis of neuroimaging, transcranial magnetic stimulation and brain-damaged patients studies. NeuroImage: Clinical, 24, 102053. 10.1016/j.nicl.2019.102053

Fedorenko, E., Hsieh, P., Nieto-Castañón, A., Whitfield-Gabrieli, S., & Kanwisher, N. (2010). New method for FMRI investigations of language: defining ROIS functionally in individual subjects. Journal of Neurophysiology, 104(2), 1177–1194. 10.1152/jn.00032.2010

Fischl, B. (2012). FreeSurfer. NeuroImage, 62(2), 774–781. 10.1016/j.neuroimage.2012.01.021

Fornaciai, M., Brannon, E. M., Woldorff, M. G., & Park, J. (2017). Numerosity processing in early visual cortex. NeuroImage, 157, 429–438. 10.1016/j.neuroimage.2017.05.069

Fornaciai, M., & Park, J. (2018). Early numerosity encoding in visual cortex is not sufficient for the representation of numerical magnitude. Journal of Cognitive Neuroscience, 30(12), 1788–1802. 10.1162/jocn_a_01320

Frey, M., Nau, M., & Doeller, C. F. (2021). Magnetic resonance-based eye tracking using deep neural networks. Nature Neuroscience, 24(12), 1772–1779. 10.1038/s41593-021-00947-w

Gallistel, C. R., & Gelman, R. (1992). Preverbal and verbal counting and computation. Cognition, 44(1–2), 43–74. 10.1016/0010-0277(92)90050-r

Gebuis, T., Gevers, W., & Kadosh, R. C. (2014). Topographic representation of high-level cognition: numerosity or sensory processing? Trends in Cognitive Sciences, 18(1), 1–3. 10.1016/j.tics.2013.10.002

Grotheer, M., Jeska, B., & Grill-Spector, K. (2018). A preference for mathematical processing outweighs the selectivity for Arabic numbers in the inferior temporal gyrus. NeuroImage, 175, 188–200. 10.1016/j.neuroimage.2018.03.064

Harvey, B. M., Klein, B. P., Petridou, N., & Dumoulin, S. O. (2013). Topographic Representation of Numerosity in the Human Parietal Cortex. Science, 341(6150), 1123–1126. 10.1126/science.1239052

Harvey, B. M., & Dumoulin, S. O. (2017). Can responses to basic non-numerical visual features explain neural numerosity responses? NeuroImage, 149, 200–209. 10.1016/j.neuroimage.2017.02.012

Harvey, B. M., & Dumoulin, S. O. (2017). A network of topographic numerosity maps in human association cortex. Nature Human Behaviour, 1(2). 10.1038/s41562-016-0036

Halberda, J., Mazzocco, M. M., & Feigenson, L. (2008). Individual differences in non-verbal number acuity correlate with maths achievement. Nature, 455(7213), 665–668. 10.1038/nature07246

Hyde, D. C., Boas, D. A., Blair, C., & Carey, S. (2010). Near-infrared spectroscopy shows right parietal specialization for number in pre-verbal infants. NeuroImage, 53(2), 647–652. 10.1016/j.neuroimage.2010.06.030

Izard, V., Dehaene-Lambertz, G., & Dehaene, S. (2008). Distinct cerebral pathways for object identity and number in human infants. PLOS Biology, 6(2), e11. 10.1371/journal.pbio.0060011

Julian, J. B., Fedorenko, E., Webster, J., & Kanwisher, N. (2012). An algorithmic method for functionally defining regions of interest in the ventral visual pathway. NeuroImage, 60(4), 2357–2364. 10.1016/j.neuroimage.2012.02.055

Kasper, L., Bollmann, S., Diaconescu, A. O., Hutton, C., Heinzle, J., Iglesias, S., Hauser, T. U., Sebold, M., Manjaly, Z., Pruessmann, K. P., & Stephan, K. Ε. (2017). The PhysIO toolbox for modeling physiological noise in FMRI data. Journal of Neuroscience Methods, 276, 56–72. 10.1016/j.jneumeth.2016.10.019

Kay, K., & Yeatman, J. D. (2017). Bottom-up and top-down computations in word- and face-selective cortex. eLife, 6. 10.7554/elife.22341

Kersey, A. J., Wakim, K., Li, R., & Cantlon, J. F. (2019). Developing, mature, and unique functions of the child’s brain in reading and mathematics. Developmental Cognitive Neuroscience, 39, 100684. 10.1016/j.dcn.2019.100684

Kim, G., Jang, J., Baek, S., Song, M., & Paik, S. B. (2021). Visual number sense in untrained deep neural networks. Science Advances, 7(1), eabd6127. 10.1126/sciadv.abd6127

Khaligh-Razavi, S., & Kriegeskorte, N. (2014). Deep Supervised, but Not Unsupervised, Models May Explain IT Cortical Representation. PLOS Computational Biology, 10(11), e1003915. 10.1371/journal.pcbi.1003915

Khaligh-Razavi, S., Cichy, R. M., Pantazis, D., & Oliva, A. (2018). Tracking the spatiotemporal neural dynamics of real-world object size and animacy in the human brain. Journal of Cognitive Neuroscience, 30(11), 1559–1576. 10.1162/jocn_a_01290

Khosla, M., & Williams, A. H. (2023). Soft Matching Distance: A metric on neural representations that captures single-neuron tuning. arXiv (Cornell University). 10.48550/arxiv.2311.09466

Knops, A., Piazza, M., Sengupta, R., Eger, E., & Melcher, D. (2014). A shared, flexible neural map architecture reflects capacity limits in both visual Short-Term memory and enumeration. Journal of Neuroscience, 34(30), 9857–9866. 10.1523/jneurosci.2758-13.2014

Kobylkov, D., Mayer, U., Zanon, M., & Vallortigara, G. (2022). Number neurons in the nidopallium of young domestic chicks. Proceedings of the National Academy of Sciences of the United States of America, 119(32). 10.1073/pnas.2201039119

Kriegeskorte, N., Mur, M., & Bandettini, P. A. (2008). Representational similarity analysis – connecting the branches of systems neuroscience. Frontiers in Systems Neuroscience. 10.3389/neuro.06.004.2008

Kriegeskorte, N., & Kievit, R. A. (2013). Representational geometry: integrating cognition, computation, and the brain. Trends in Cognitive Sciences, 17(8), 401–412. 10.1016/j.tics.2013.06.007

Kriegeskorte, N., & Wei, X. (2021). Neural tuning and representational geometry. Nature Reviews. Neuroscience, 22(11), 703-718. 10.1038/s41583-021-00502-3

Kruskal, J. B. (1964). Multidimensional scaling by optimizing goodness of fit to a nonmetric hypothesis. Psychometrika, 29(1), 1–27. 10.1007/bf02289565

Kutter, E. F., Bostroem, J., Elger, C. E., Mormann, F., & Nieder, A. (2018). Single neurons in the human brain encode numbers. Neuron, 100(3), 753–761.e4. 10.1016/j.neuron.2018.08.036

Kutter, E. F., Boström, J., Elger, C. E., Nieder, A., & Mormann, F. (2022). Neuronal codes for arithmetic rule processing in the human brain. Current Biology, 32(6), 1275–1284.e4. 10.1016/j.cub.2022.01.054

Kutter, E. F., Dehnen, G., Borger, V., Surges, R., Mormann, F., & Nieder, A. (2023). Distinct neuronal representation of small and large numbers in the human medial temporal lobe. Nature Human Behaviour. 10.1038/s41562-023-01709-3\

Lasne, G., Piazza, M., Dehaene, S., Kleinschmidt, A., & Eger, E. (2019). Discriminability of numerosity-evoked fMRI activity patterns in human intra-parietal cortex reflects behavioral numerical acuity. Cortex, 114, 90–101. 10.1016/j.cortex.2018.03.008

Lemer, C., Dehaene, S., Spelke, E. S., & Cohen, L. (2003). Approximate quantities and exact number words: dissociable systems. Neuropsychologia, 41(14), 1942–1958. 10.1016/s0028-3932(03)00123-4

Luyckx, F., Nili, H., Spitzer, B., & Summerfield, C. (2019). Neural structure mapping in human probabilistic reward learning. eLife, 8. 10.7554/elife.42816

Merten, K., & Nieder, A. (2009). Compressed scaling of abstract numerosity representations in adult humans and monkeys. Journal of Cognitive Neuroscience, 21(2), 333–346. 10.1162/jocn.2008.21032

Mistry, P. K., Strock, A., Liu, R., Young, G. B., & Menon, V. (2023). Learning-induced reorganization of number neurons and emergence of numerical representations in a biologically inspired neural network. Nature Communications, 14(1). 10.1038/s41467-023-39548-5

Mitchell, T. M., Hutchinson, R., Niculescu, R. S., Pereira, F., Wang, X., Just, M. A., & Newman, S. D. (2004). Learning to Decode Cognitive States from Brain Images. Machine Learning, 57(1/2), 145–175. 10.1023/b:mach.0000035475.85309.1b

Moeller, S., Yacoub, E., Olman, C. A., Auerbach, E. J., Strupp, J., Harel, N., & Uğurbil, K. (2010). Multiband multislice GE-EPI at 7 tesla, with 16-fold acceleration using partial parallel imaging with application to high spatial and temporal whole-brain fMRI. Magnetic Resonance in Medicine, 63(5), 1144–1153. 10.1002/mrm.22361

Moscoso, P. a. M., Greenlee, M. W., Anobile, G., Arrighi, R., Burr, D., & Castaldi, E. (2021). Groupitizing modifies neural coding of numerosity. Human Brain Mapping, 43(3), 915–928. 10.1002/hbm.25694

Nau, M., Schindler, A., & Bartels, A. (2018). Real-motion signals in human early visual cortex. NeuroImage, 175, 379–387. 10.1016/j.neuroimage.2018.04.012

Nau, M., Schröder, T., Bellmund, J. L. S., & Doeller, C. F. (2018). Hexadirectional coding of visual space in human entorhinal cortex. Nature Neuroscience, 21(2), 188–190. 10.1038/s41593-017-0050-8

Nasr, K., Viswanathan, P., & Nieder, A. (2019). Number detectors spontaneously emerge in a deep neural network designed for visual object recognition. Science Advances, 5(5), eaav7903. 10.1126/sciadv.aav7903

Nelli, S., Braun, L., Dumbalska, T., Saxe, A. M., & Summerfield, C. (2023). Neural knowledge assembly in humans and neural networks. Neuron, 111(9), 1504–1516.e9. 10.1016/j.neuron.2023.02.014

Nieder, A., & Miller, E. K. (2003). Coding of Cognitive Magnitude. Neuron, 37(1), 149–157. 10.1016/s0896-6273(02)01144-3

Nieder, A., & Miller, E. K. (2004). A parieto-frontal network for visual numerical information in the monkey. Proceedings of the National Academy of Sciences of the United States of America, 101(19), 7457–7462. 10.1073/pnas.0402239101

Nieder, A. (2020). The Adaptive Value of Numerical Competence. Trends in Ecology & Evolution, 35(7), 605–617. 10.1016/j.tree.2020.02.009

Nieto-Castañón, A., & Fedorenko, E. (2012). Subject-specific functional localizers increase sensitivity and functional resolution of multi-subject analyses. NeuroImage, 63(3), 1646–1669. 10.1016/j.neuroimage.2012.06.065

Nili, H., Wingfield, C., Walther, A., Su, L., Marslen-Wilson, W. D., & Kriegeskorte, N. (2014). A toolbox for representational similarity analysis. PLOS Computational Biology, 10(4), e1003553. 10.1371/journal.pcbi.1003553

Oosterhof, N. N., Wiestler, T., Downing, P. E., & Diedrichsen, J. (2011). A comparison of volume-based and surface-based multi-voxel pattern analysis. NeuroImage, 56(2), 593–600. 10.1016/j.neuroimage.2010.04.270

Oosterhof, N. N., Connolly, A. C., & Haxby, J. V. (2016). COSMOMVPA: Multi-Modal Multivariate Pattern Analysis of Neuroimaging data in Matlab/GNU Octave. Frontiers in Neuroinformatics, 10. 10.3389/fninf.2016.00027

Piantadosi, S. T., & Cantlon, J. F. (2017). True numerical cognition in the wild. Psychological Science, 28(4), 462–469. 10.1177/0956797616686862

Piazza, M., Izard, V., Pinel, P., le Bihan, D., & Dehaene, S. (2004). Tuning Curves for Approximate Numerosity in the Human Intraparietal Sulcus. Neuron, 44(3), 547–555. 10.1016/j.neuron.2004.10.014

Piazza, M., Pinel, P., le Bihan, D., & Dehaene, S. (2007). A Magnitude Code Common to Numerosities and Number Symbols in Human Intraparietal Cortex. Neuron, 53(2), 293–305. 10.1016/j.neuron.2006.11.022

Piazza, M. (2010). Neurocognitive start-up tools for symbolic number representations. Trends in Cognitive Sciences, 14(12), 542–551. 10.1016/j.tics.2010.09.008

Piazza, M., & Eger, E. (2016). Neural foundations and functional specificity of number representations. Neuropsychologia, 83, 257–273. 10.1016/j.neuropsychologia.2015.09.025

Polti, I., Nau, M., Kaplan, R., Van Wassenhove, V., & Doeller, C. F. (2022). Rapid encoding of task regularities in the human hippocampus guides sensorimotor timing. eLife, 11. 10.7554/elife.79027

Paul, J., Van Ackooij, M., Cate, T. C. T., & Harvey, B. M. (2022). Numerosity tuning in human association cortices and local image contrast representations in early visual cortex. Nature Communications, 13(1). 10.1038/s41467-022-29030-z

Revkin, S. K., Piazza, M., Izard, V., Cohen, L., & Dehaene, S. (2008). Does subitizing reflect numerical estimation? Psychological Science, 19(6), 607–614. 10.1111/j.1467-9280.2008.02130.x

Rosenke, M., Van Hoof, R., Van Den Hurk, J., Grill-Spector, K., & Goebel, R. (2020). A Probabilistic Functional Atlas of Human Occipito-Temporal Visual Cortex. Cerebral Cortex, 31(1), 603–619. 10.1093/cercor/bhaa246

Saad, Z. S., Reynolds, R. C., Argall, B., Japee, S., & Cox, R. W. (2005). SUMA: An interface for surface-based intra- and inter-subject analysis with AFNI. IEEE International Symposium on Biomedical Imaging: From Nano to Macro. 10.1109/isbi.2004.1398837

Sani, I., Stemmann, H., Caron, B., Bullock, D., Stemmler, T., Fahle, M., Pestilli, F., & Freiwald, W. A. (2021). The human endogenous attentional control network includes a ventro-temporal cortical node. Nature Communications, 12(1). 10.1038/s41467-020-20583-5

Scott, T. L., & Perrachione, T. K. (2019). Common cortical architectures for phonological working memory identified in individual brains. NeuroImage, 202, 116096. 10.1016/j.neuroimage.2019.116096

Smith, S. M., & Nichols, T. E. (2009). Threshold-free cluster enhancement: Addressing problems of smoothing, threshold dependence and localisation in cluster inference. NeuroImage, 44(1), 83–98. 10.1016/j.neuroimage.2008.03.061

Stemmann, H., & Freiwald, W. A. (2019). Evidence for an attentional priority map in inferotemporal cortex. Proceedings of the National Academy of Sciences of the United States of America, 116(47), 23797–23805. 10.1073/pnas.1821866116

Verma, B. K., & Sengupta, R. (2023). Emergence of behavioral phenomena and adaptation effects in human numerosity decoder using recurrent neural networks. Scientific Reports, 13(1). 10.1038/s41598-023-44535-3

Viswanathan, P., Stein, A. M., & Nieder, A. (2024). Sequential neuronal processing of number values, abstract decision, and action in the primate prefrontal cortex. PLOS Biology, 22(2), e3002520. 10.1371/journal.pbio.3002520

Wagener, L., Loconsole, M., Ditz, H. M., & Nieder, A. (2018). Neurons in the Endbrain of Numerically Naive Crows Spontaneously Encode Visual Numerosity. Current Biology, 28(7), 1090–1094.e4. 10.1016/j.cub.2018.02.023

Wang, L., Mruczek, R. E. B., Arcaro, M., & Kästner, S. (2014). Probabilistic maps of visual topography in human cortex. Cerebral Cortex, 25(10), 3911–3931. 10.1093/cercor/bhu277

Warrington, E. K., & James, M. (1967). Tachistoscopic number estimation in patients with unilateral cerebral lesions. Journal of Neurology, Neurosurgery, and Psychiatry, 30(5), 468–474. 10.1136/jnnp.30.5.468

Yeo, D. J., Wilkey, E. D., & Price, G. R. (2017). The search for the number form area: A functional neuroimaging meta-analysis. Neuroscience & Biobehavioral Reviews, 78, 145–160. 10.1016/j.neubiorev.2017.04.027

