## Supplementary File for "Distinct neural representational geometries of numerosity in early visual and association regions of dorsal and ventral streams"

### Supplementary material

#### S1. Further exploration of the neural representational geometries in early and associative regions of both streams

In order to test whether we could capture additional variance and reveal potential additional hidden structures that might not have been apparent in the first two dimensions, we also examined the third dimension of the MDS analysis of the neural RDMs in all different retinotopically organized regions along both the ventral and the dorsal stream. The results indicate that:

1. Adding the third dimension did not significantly increase the variance already explained by the first two (see figure S1). We computed the variance explained by dividing the square of each eigenvalue by the sum of the squares of all eigenvalues as generated by the function *cmdscale* in MATLAB.

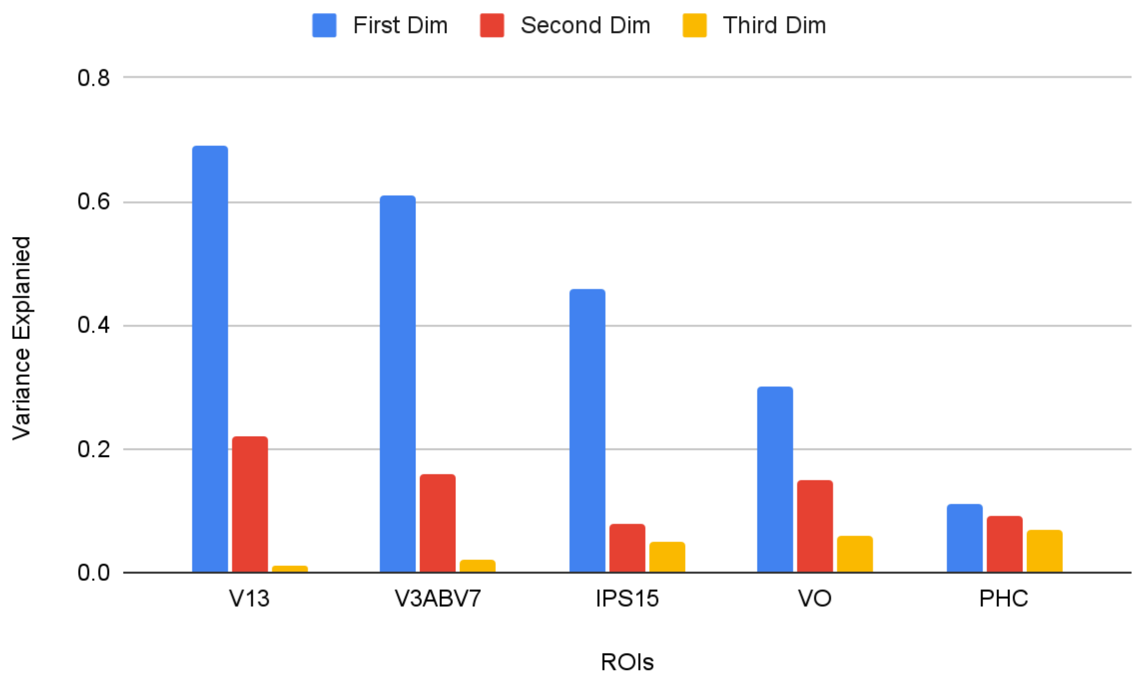


Figure S1: Explained variance by the first three dimensions obtained from classical multidimensional scaling (MDS) applied to data from V13, V3A/V7, IPS15, VO, and PHC.

2. The third dimension did not add novel insight on the geometry of the neural representation of stimuli (see Figure S2, where the results of the MDS is visualized in 3D, with average values in red, and 2D projections mapped in blue. It is clear that in early visual regions as well as in VO along the ventral stream, all three dimensions are a straight line, while progressing along the dorsal stream all tend to display curvature.

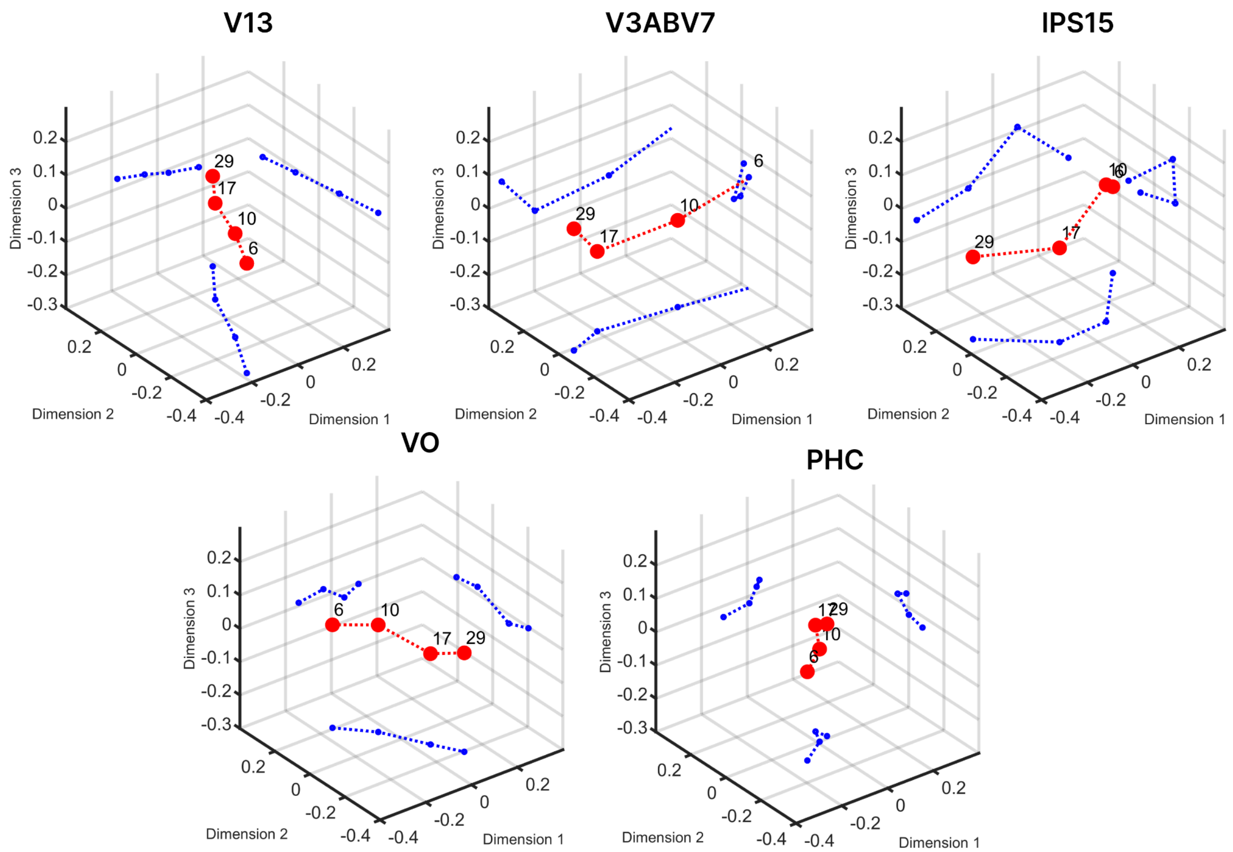


Figure S2: Three-dimensional visualization of the representational geometry obtained from classical multidimensional scaling (MDS) applied to RDMs from V13, V3A/V7, IPS15, VO, and PHC.

#### S2. Simulations

In order to verify whether the linear vs. curved structures that we observed in the neural RDMs could result from two different underlying neuronal tuning schemes, we performed some simulations. The first step in these simulations consisted in assuming that different voxels are tuned to different numbers, considering two different coding schemes: monotonic and tuned. In both cases we simulated 4,200 voxels and we started by assuming that the same proportion of voxels would code each numerosity and that the widths of the tuning would be relatively small (we considered three SDs separately: 0.3, 0.4, and 0.5).

In the case of monotonic response profiles, we set up artificial voxels exhibiting either a square-root or logarithmic response function to 4, 6, 10, 17, 29, and 50. This setup modeled a scenario in which voxel activity systematically increases with numerosity.

In the case of numerosity-tuned voxels with Gaussian response functions, we modelled numerosity-tuned voxels with Gaussian response functions using both the continuous number range from 1 to 40 and the specific number ranges of 4, 6, 10, 17, 29, and 50.

As a second step of our simulations we computed the pattern dissimilarity between individual conditions using Pearson correlation distance as the distance measure, as we did in the original RSA analyses based on real BOLD signal data. On the basis of these data we generated two distinct sets of representational dissimilarity matrices (RDMs), from which we computed the multidimensional scaling (MDS) and visualized the response. The resulting representational dissimilarity matrices from these simulations are depicted in Figure S3, and the corresponding multidimensional scaling (MDS) visualization is shown in Figure S4. Given that the observed results were in line with the expectations, we then checked if they would hold using more biologically plausible conditions. In particular, on the bases of Cai et al., 2021, we consider two properties of the fMRI tuning preference to numerosity:

1) The voxels tuned to the different numerosities decrease with numerosity (see Cai et al., 2021, Fig. 3). To simulate this, we allocated half of the voxels to the number range 7–16, approximately 57% of the remaining half to 16–32, and the rest to 32–64. We repeated the analysis for number ranges: 4, 6, 10, 17, 29, and

50 (See Figure S5). The results were substantially identical to those derived from a fixed distribution of tuning preferences.

2) Second, the tuning curves for numerosity in fMRI are considerably broader than those used in our initial simulation. For example, the Full Width at Half Maximum (FWHM) for small numerosities such as two was approximately 9 (see Cai et al., 2021, Fig. 1). To assess the effect of broader tuning, we repeated the analysis using a larger standard deviation (SD) for the tuning curves. As in the previous condition, we simulated the number ranges 4, 6, 10, 17, 29, and 50, while also incorporating the biologically inspired constraint that the number of tuned voxels decreases with numerosity. However, under this condition, the curved structure in the MDS solution disappeared (see Figure S6 part A).
3) Finally, we also considered another empirical finding from Cai et al. (2021): the tuning width increases linearly with numerosity (see their Fig. 4). Accordingly, in addition to incorporating the two biological constraints described above—the decreasing number of tuned voxels with increasing numerosity and the broader overall tuning width—we conducted a simulation in which the standard deviation (SD) of the tuning curves increased proportionally with numerosity. As in the previous conditions, we used the number ranges 4, 6, 10, 17, 29, and 50. This allowed us to examine the combined effect of all biologically inspired constraints on the resulting representational structure. However, under this condition again, the curved structure in the MDS solution disappeared (see Figure S6, part B).

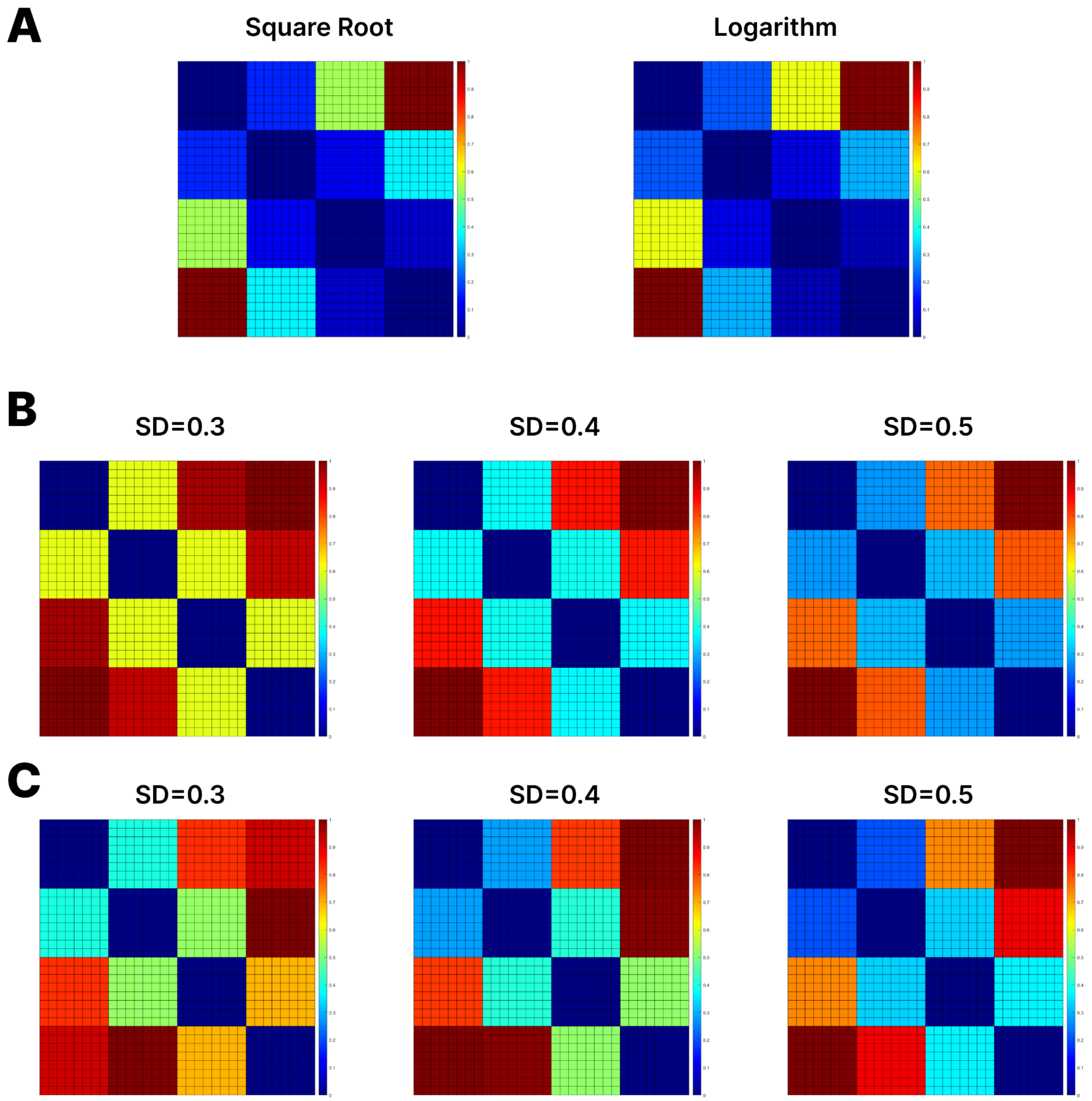


Figure S3. (A) RDMs resulted from monotonic simulation using square root and logarithmic functions for number ranges of 4, 6, 10, 17, 29, and 50. (B) Simulation with Gaussian functions of varying widths (0.3, 0.4, 0.5) for number ranges of 4, 6, 10, 17, 29, and 50. (C) Simulation with Gaussian functions of varying widths (0.3, 0.4, 0.5) for a continuous number range from 1 to 40, including all intermediate values.

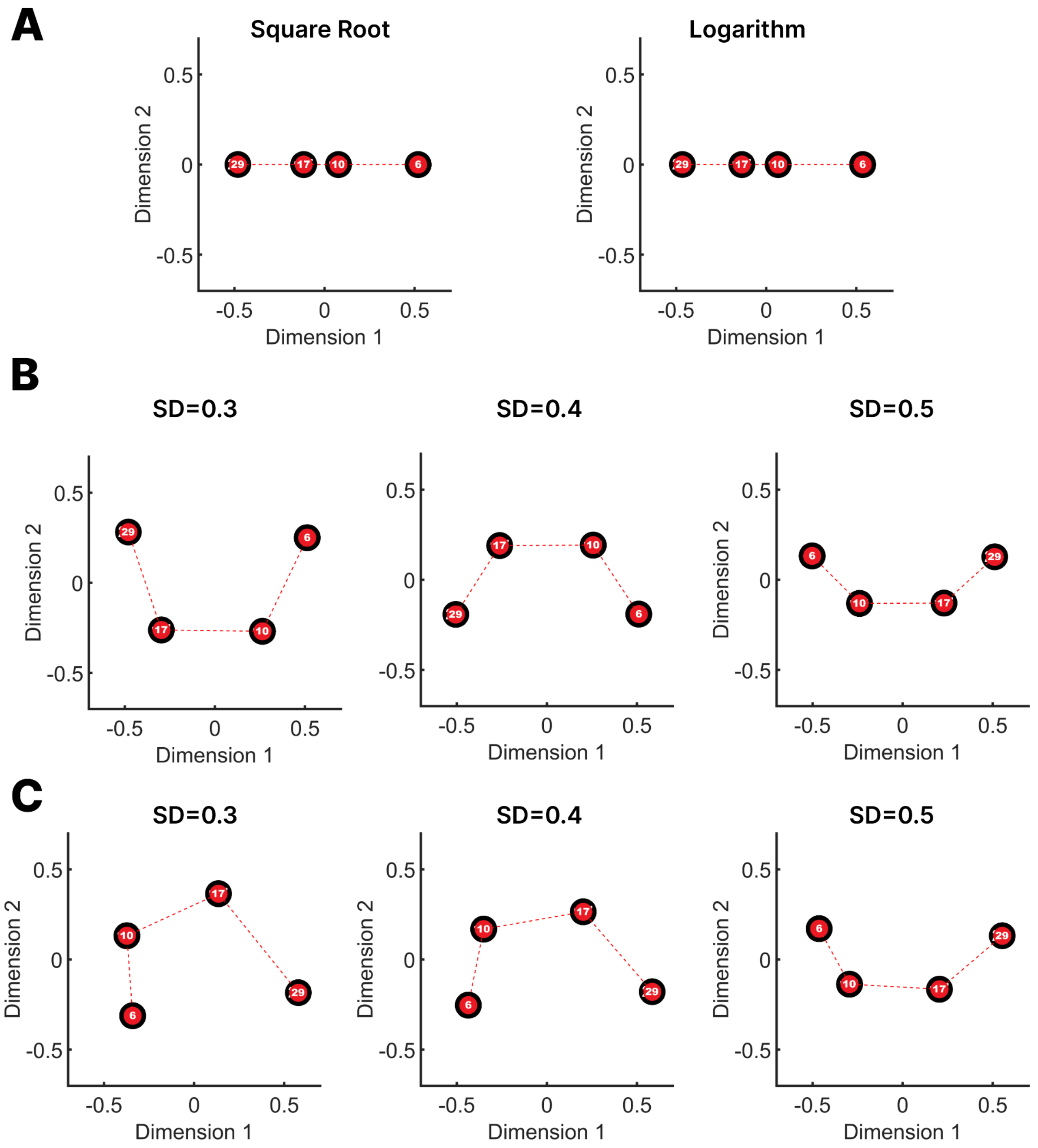


Figure S4. (A) MDS resulted from monotonic simulation using square root and logarithmic functions for number ranges of 4, 6, 10, 17, 29, and 50. (B) Simulation with Gaussian functions of varying widths (0.3, 0.4, 0.5) for number ranges of 4, 6, 10, 17, 29, and 50. (C) Simulation with Gaussian functions of varying widths (0.3, 0.4, 0.5) for a continuous number range from 1 to 40, including all intermediate values.

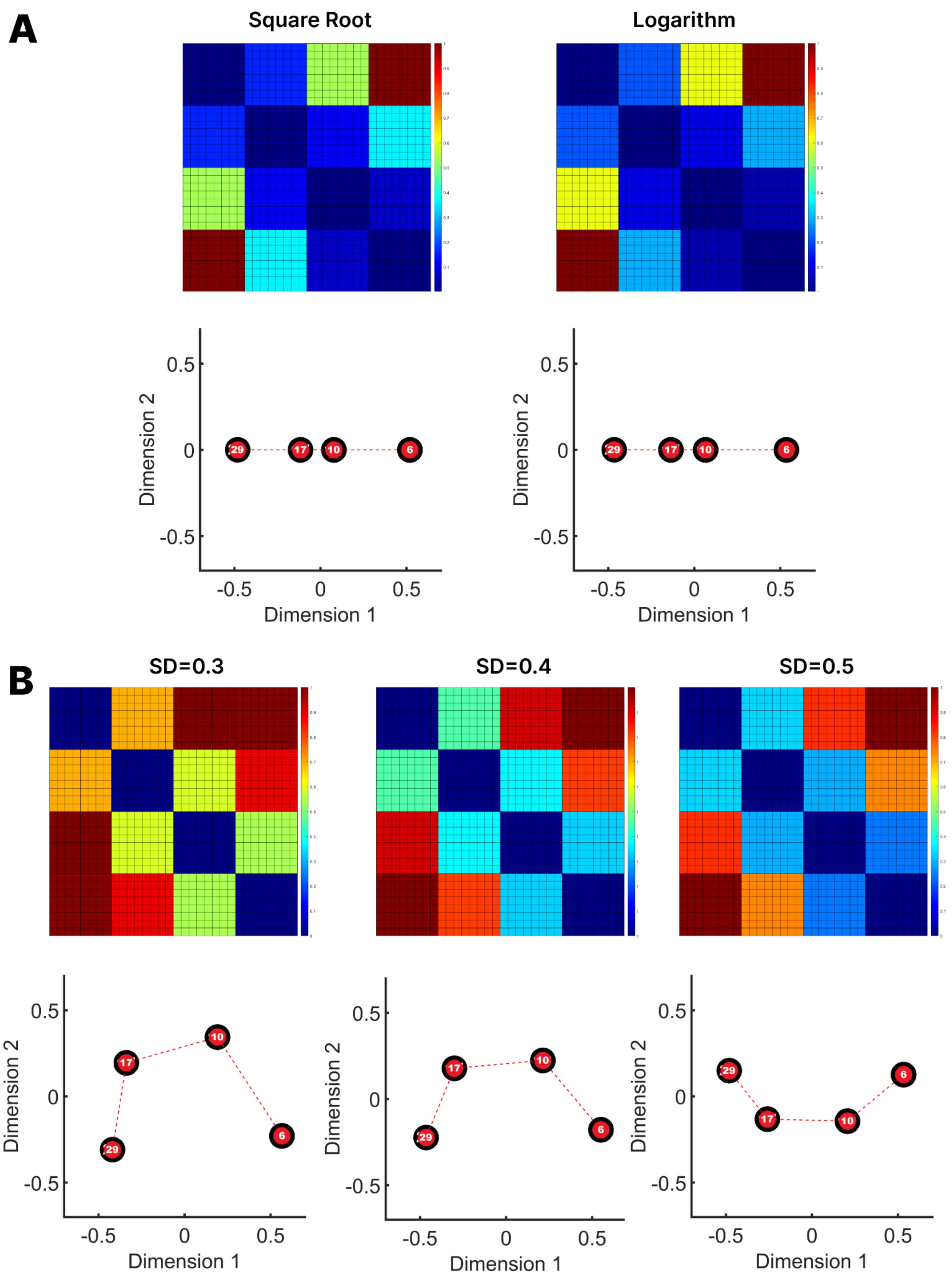


Figure S5. (A) RDM and MDS results from monotonic simulations using square root and logarithmic functions for number ranges of 4, 6, 10, 17, 29, and 50. (B) Simulations using Gaussian tuning functions of varying widths (0.3, 0.4, 0.5) for the same number ranges. In both cases, different proportions of voxels were allocated to each number range, following a biologically inspired distribution.

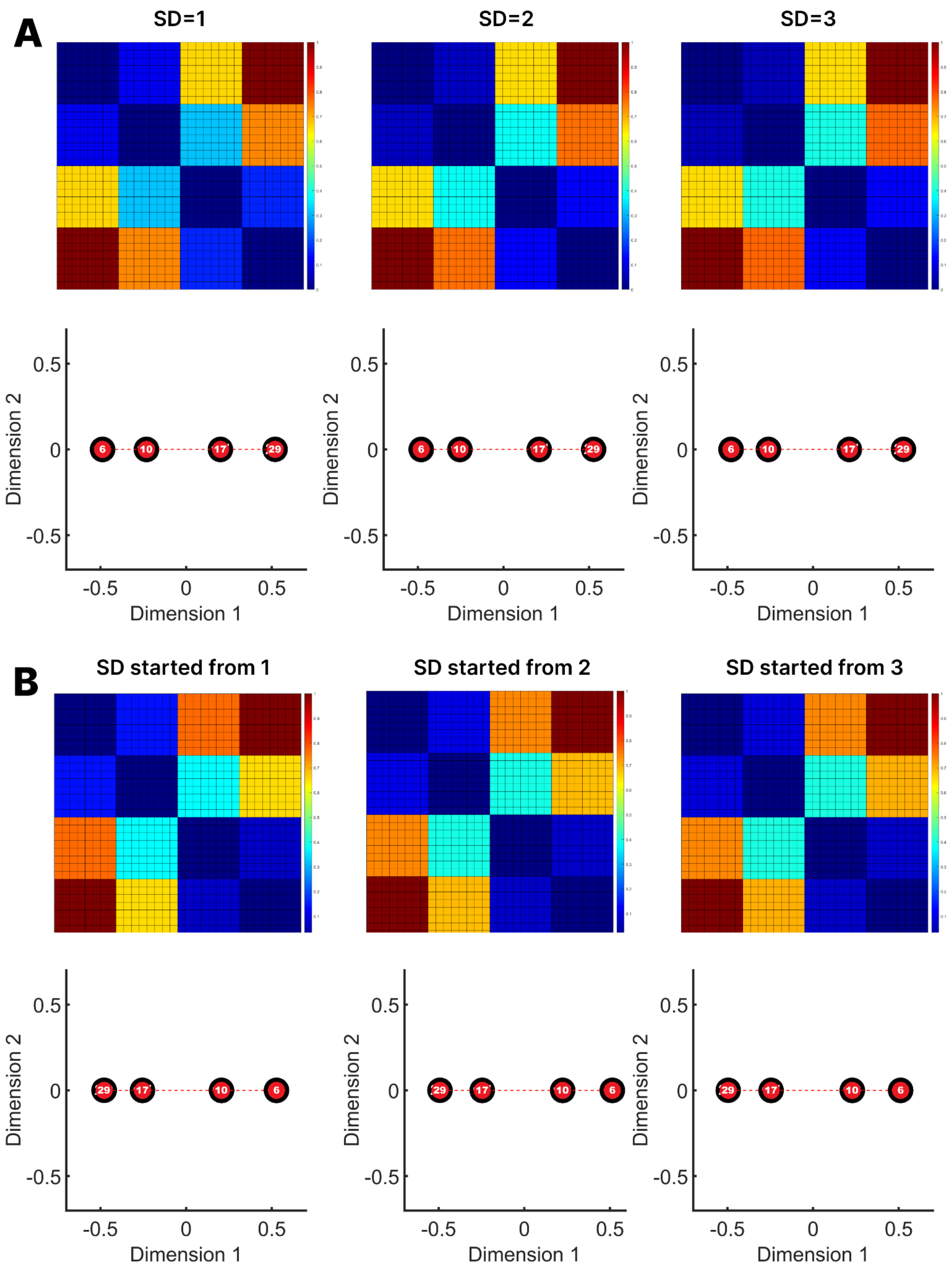


Figure S6. (A) RDM and MDS resulted from simulation with Gaussian functions of varying widths (1, 2, 3) for number ranges of 4, 6, 10, 17, 29, and 50. (B) RDM and MDS results from simulations using Gaussian tuning functions with linearly increasing standard deviations, starting at 1, 2, or 3 and increasing by 1 unit for each subsequent number. Simulations were performed for the number ranges 4, 6, 10, 17, 29, and 50. In both cases, different proportions of voxels were allocated to each number range, following a biologically inspired distribution.

Cai, Y., Hofstetter, S., Van Dijk, J., Zuiderbaan, W., Van Der Zwaag, W., Harvey, B. M., & Dumoulin, S. O. (2021). Topographic numerosity maps cover subitizing and estimation ranges. Nature Communications, 12(1). https://doi.org/10.1038/s41467-021-23785-7

De Beeck, H. P. O., Baker, C. I., DiCarlo, J. J., & Kanwisher, N. G. (2006). Discrimination training alters object representations in human extrastriate cortex. Journal of Neuroscience, 26(50), 13025–13036. https://doi.org/10.1523/jneurosci.2481-06.2006

Garrido, L., Vaziri-Pashkam, M., Nakayama, K., & Wilmer, J. (2013). The consequences of subtracting the mean pattern in fMRI multivariate correlation analyses. Frontiers in Neuroscience, 7. https://doi.org/10.3389/fnins.2013.00174

Haxby, J. V., Gobbini, M. I., Furey, M. L., Ishai, A., Schouten, J. L., & Pietrini, P. (2001). Distributed and overlapping representations of faces and objects in ventral temporal cortex. Science, 293(5539), 2425–2430. https://doi.org/10.1126/science.1063736

Sayres, R., & Grill-Spector, K. (2008). Relating Retinotopic and Object-Selective responses in human lateral occipital cortex. Journal of Neurophysiology, 100(1), 249–267. https://doi.org/10.1152/jn.01383.2007
